# An intracellular calcium sensory complex CPKs/ECA1 controls cytosolic calcium homeostasis for plant osmosensitivity

**DOI:** 10.1101/2024.10.13.617954

**Authors:** Xiaoju Liang, Yeling Zhou, Weifeng Xu, Jiansheng Liang

**Author notes:** These authors contributed equally to this work (Xiaoju Liang, Yeling Zhou). To whom correspondence may be addressed. (W. X); (J. L).

## Abstract

Intracellular Ca^2+^ controls various cellular functions and local Ca^2+^ dynamics is tightly regulated upon environmental cues. Maintaining cellular Ca^2+^ balance is essential for plant survival. Here we report a calcium-dependent protein kinases (CPKs)-mediated signaling pathway, in conjunction with the ER membrane-resident Ca^2+^-ATPase ECA1, acts influentially for cytosolic Ca^2+^ homeostasis and osmotic stress tolerance. We show that targeting cytosolic Ca^2+^ efflux via specific inhibitors or *eca1* mutation results in augmented [Ca^2+^]cyt spikes, elevated cytoplasmic ABA ([ABA]cyt) level and ultimately hypersensitive to osmotic stress. Screening of *Arabidopsis* CPKs revealed direct binding of CPK2/6/11 to ECA1. Moreover, CPK2/6/11 phosphorylate the N-terminal of ECA1 at Ser5, thereby enhancing its activity for cytosolic Ca^2+^ efflux into ER and subsequently lower [ABA]cyt. The cumulative effect of ECA1 and CPKs mutation on *Arabidopsis* plant sensitivity to osmotic stress further illustrates that CPKs/ECA1 acts an intracellular sensory module for plant stress tolerance via regulating [Ca^2+^]cyt and [ABA]cyt homeostasis.

**One-sentence summary:** CPKs/ECA1 acts an intracellular sensory module for plant osmotic stress tolerance via regulating cytosolic Ca^2+^ and ABA homeostasis.

## Introduction

Terrestrial plants have adapted themselves to a constantly changing environment via a plethora of signaling molecules. Ca^2+^, among others, represents one of the most influential molecules for it is not only a critical nutrient for plant growth and development^1, 2, 3^, but also an essential cellular signal during stress responses^4, 5, 6^. Ca^2+^ signaling typically features a transient increase in the cytosolic calcium concentration ([Ca^2+^]cyt) triggered upon numerous stresses, followed by a decay that results in Ca^2+^ spikes, thereby initiating subsequent molecular and physiological responses. Ca^2+^ signal is decoded by a diverse array of calcium-binding proteins, including calmodulin (CaM), calmodulin-like proteins (CMLs), calcium-dependent protein kinases (CDPK/CPKs), and calcineurin B-like proteins (CBLs)^7, 8^. Particularly, CPKs play significant roles in multiple stress responses through phosphorylation and interconnection with other signaling pathways^9^. For example, *Arabidopsis* CPK1/4/6/11 positively regulate salt and drought stress tolerance through regulation of ABA signaling^10, 11^.

The cytosolic Ca^2+^ homeostasis is tightly controlled by diverse calcium channels, Ca^2+^-ATPase and transporters that are ubiquitously distributed in plant membranes^12, 13, 14, 15^. With an over 10, 000 times gradience between extra- and intracellular space, plant cells constantly risk excessive cytosolic Ca^2+^ ion that eventually leads to cell damage or death. To prevent this, a potent Ca^2+^ extrusion system is engaged to allow efficient Ca^2+^ efflux against the electrochemical gradient. Typically, cellular Ca^2+^ influx is facilitated by calcium-permeable cation channels, whereas Ca^2+^-ATPase and Ca^2+^/H^+^ exchangers assist Ca^2+^ back to extra- and intracellular pools^16^. Despite the growing evidence on calcium channels or pumps for mediating [Ca^2+^]cyt influx during stress response^6, 17, 18, 19, 20, 21, 22^, mechanisms underlying cytosolic Ca^2+^ removal are less understood^23, 24^. A recent study detailed two distinct signaling pathways that converge on vacuolar Ca^2+^/H^+^ exchangers (CAXs) for balancing plant growth and immunity via regulation of cytosolic Ca^2+^ efflux^25^.

Two types of Ca^2+^-ATPases exist in plants, with the autoinhibited Ca^2+^-ATPases (ACA) located at both the plasma membrane (PM) and endomembrane and the ER-type Ca^2+^-ATPases (ECA) mostly resided on the endomembrane^26^. Previous studies have implicated plant Ca^2+^-ATPases in pollination and pollen development^27, 28^, early seedling development^29^, pathogen-induced cell death^30, 31, 32^. Notably, ECA1, an ER-type Ca^2+^-ATPase, is involved in stress tolerance and plant root responses to hydrotropism via maintaining cellular Ca^2+^ homeostasis^33, 34, 35^. These findings substantiate the physiological significance of plant Ca^2+^-ATPases, but how they act in controlling [Ca^2+^]cyt efflux and subsequent plant tolerance is not clear. In this study, we identified an intracellular calcium sensory complex that acts fundamentally in maintaining [Ca^2+^]cyt and ABAcyto homeostasis. Our findings highlight the intricacy of cellular Ca^2+^ oscillations for plant to balance the stress tolerance and the growth.

## Results

### Augmentation of [Ca^2+^]cyt spikes aggravates osmosensitive root growth in an ABA-dependent manner

Being a vital second messenger in plants, [Ca^2+^]cyt has to be kept at a critically low level to avoid hazardous effect (Leitão et al., 2019). To test how perturbations in [Ca^2+^]cyt affect plant osmosensitivity, we applied two calcium ion transporter inhibitors that specifically target PM-resided Ca^2+^-ATPases (LaCl3)^36^ and the ER membrane Ca^2+^-ATPase ECA1 (cyclopiazonic acid, CPA)^34, 37^, respectively, and examined root growth under a series of mannitol treatments. Exogenous mannitol treatments led to root growth inhibition in a concentration-dependent manner (Fig. 1A, B). Whilst co-treatment of plants with LaCl3 resulted in similar root growth as those treated with mannitol alone, the presence of CPA notably shortened root length (Fig. 1A, B). Monitor of [Ca^2+^]cyt concentration using Ca^2+^ sensor lines^38^ revealed that mannitol treatment evoked a rapid and transient increase in [Ca^2+^]cyt, which was returned to basal levels around 50 s after treatment (Fig. 1C, D). Moreover, pretreatment with LaCl3 significantly diminished [Ca^2+^]cyt spikes triggered upon osmotic stress, whereas addition of CPA elicited a surprisingly elevated and prolonged [Ca^2+^]cyt surge (Fig. 1C, D). These results support that sustained [Ca^2+^]cyt elevation leads to plant hypersensitivity and other signals may override Ca^2+^ for normal response to osmotic stimulus.

**Fig. 1.**
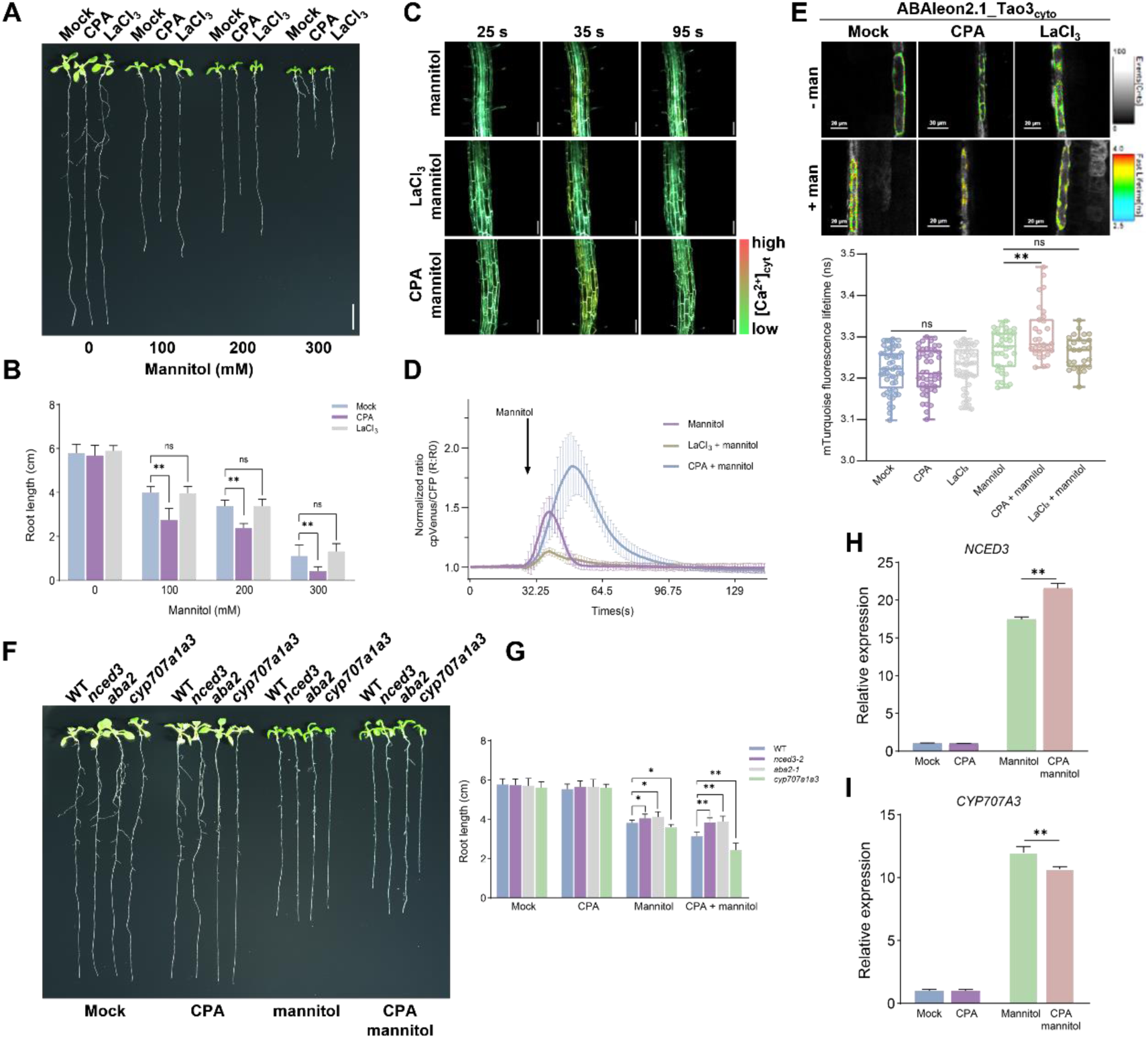
| CPA magnifies [Ca^2+^]_cyt_ spikes and osmosensitivity depending on cytosolic ABA homeostasis. **A, B** The representative image (**A**) and root length (**B**) of *Arabidopsis* Col-0 seedlings treated with mannitol and calcium inhibitors. Four-day-old seedlings were transferred to ½ MS medium containing 0, 100, 200, 300, 400 mM mannitol in the presence of CPA (10 μM) or LaCl_3_ (50 μM) and grown for 7 days before imaging analysis. Data are means ± SD, n = 15. Statistics were performed using one-way ANOVA followed by Dunnett’s multiple comparisons test within the same mannitol concentration treatment (\*\**P* < 0.01; ns, not significant). **C, D** Representative images (**C**) and normalized ratio values (ΔR:R0) of [Ca^2+^]_cyt_ changes (**D**) in response to mannitol and inhibitors treatment. Six-day-old Arabidopsis roots expressing the Ca^2+^ sensor YC3.6 were pretreated with LaCl_3_ (50 μM) or CPA (25 μM) for 15 min prior to mannitol treatment (400 mM). The pseudo-color scale bar along the images shows colors from green to red, representing the [Ca^2+^]_cyt_ level from low to high. The arrow in (**D**) indicates the start point of mannitol application. The ratio graphs represent the average ±SD, n = 3. **E** Representation FLIM images and data showing the changes of cytosolic ABA level in response to mannitol in the presence or absence of calcium inhibitors. Seven-day-old Arabidopsis roots expressing the ABA sensor ABAleon2.1_Tao3_cyt_ were pretreated with LaCl_3_ (50 μM) or CPA (5 μM) before exposed to mannitol (300 mM) for 4 h. Fluorescence lifetime data are shown as box plot with original data points. Statistics were performed using one-way ANOVA followed by Dunnett’s multiple comparisons test between treatments (\*\**P* < 0.01; ns, not significant). **F, G** The representative image (**F**) and root length (**G**) of Arabidopsis wild-type Col-0 and ABA mutants in response to mannitol and CPA. Four-day-old seedlings were treated with CPA (5 μM), mannitol (200 mM), or mannitol + CPA for 7 days before analysis. Statistics were performed using one-way ANOVA followed by Dunnett’s multiple comparisons test between genotypes within the same treatment (\**P* < 0.05; \*\**P* < 0.01; ns, not significant). **H, I** Relative expression of *NCED3* (**H**) and *CYP707A3* (**I**) upon mannitol and CPA treatments. Eight-day-old seedlings were transferred onto ½ MS medium ± 10 μM CPA supplemented with or without 200 mM mannitol for 1 hour. Values are means ± SD, n = 3. Statistics were performed using Students’ *t* test (\*\**P* < 0.01).

Osmotic stress triggers a swift accumulation of ABA within hours^39^. To investigate if perturbations in [Ca^2+^]cyt spikes impact ABA levels, we tested the effect of calcium inhibitors on cytosolic and ER ABA levels using our previously developed ABA sensor lines^40, 41^. *Arabidopsis* seedlings co-treated with CPA and mannitol accumulated a notably higher level of cytosolic ABA compared with those treated mannitol alone or co-treated with LaCl3 (Fig. 1E). Similarly enhanced ABAcyto levels triggered upon CPA and mannitol treatments were observed in transient tobacco expression system (Supplementary Fig. 1). These results suggest that ABA acts as a downstream signal of Ca^2+^ for plant osmosensitivity. Furthermore, the CPA-triggered hypersensitivity to osmotic stimulus was abolished in ABA biosynthetic mutants *nced3-2* and *aba2-1*, whereas it was aggravated in ABA metabolic mutant *cyp707a1a3* that contains more endogenous ABA (Kushiro et al., 2004) (Fig. 1F). In accordance, gene expression analysis showed that the expression level of the rate-limiting *NCED3* was significantly enhanced under CPA and mannitol cotreatment, whereas the ABA catabolic gene *CYP707A3* was reduced in comparison to those under mannitol treatment alone (Fig. 1H, I). These findings together point to a fact that excess of [Ca^2+^]cyt causes hypersensitive root responses to osmotic stress via alteration of cytosolic ABA homeostasis.

### ECA1 participates in the regulation of osmosensitivity via changing the [Ca^2+^]cyt and [ABA]cyt levels

Given that CPA specifically targets and inhibits ECA1 activity^37^, we expected a role of ECA1 in *Arabidopsis* root responses to osmotic stimulus. To test that, we obtained a T-DNA insertion mutant *eca1-1* and generated *eca1-2* using the clustered regularly interspaced short palindromic repeat (CRISPR)/CRISPR-associated nuclease 9 (Cas9) system^42^ (Supplementary Fig. 2). Besides, we generated transgenic lines overexpressing *ECA1* driven by the Cauliflower mosaic virus (CaMV35S) promoter (Supplementary Fig. 2B). Both *eca1-1* and *eca1-2* mutants displayed shorter roots compared with Col-0 upon mannitol treatments, whereas the complementary lines that obtained by crossing the overexpressing lines with the *eca1-1* mutant showed similar osmotic-trigged root growth inhibition as wild-type (WT) Col-0 (Fig. 2A, B). These results support an important role of ECA1 in root elongation under osmotic stress.

**Fig. 2.**
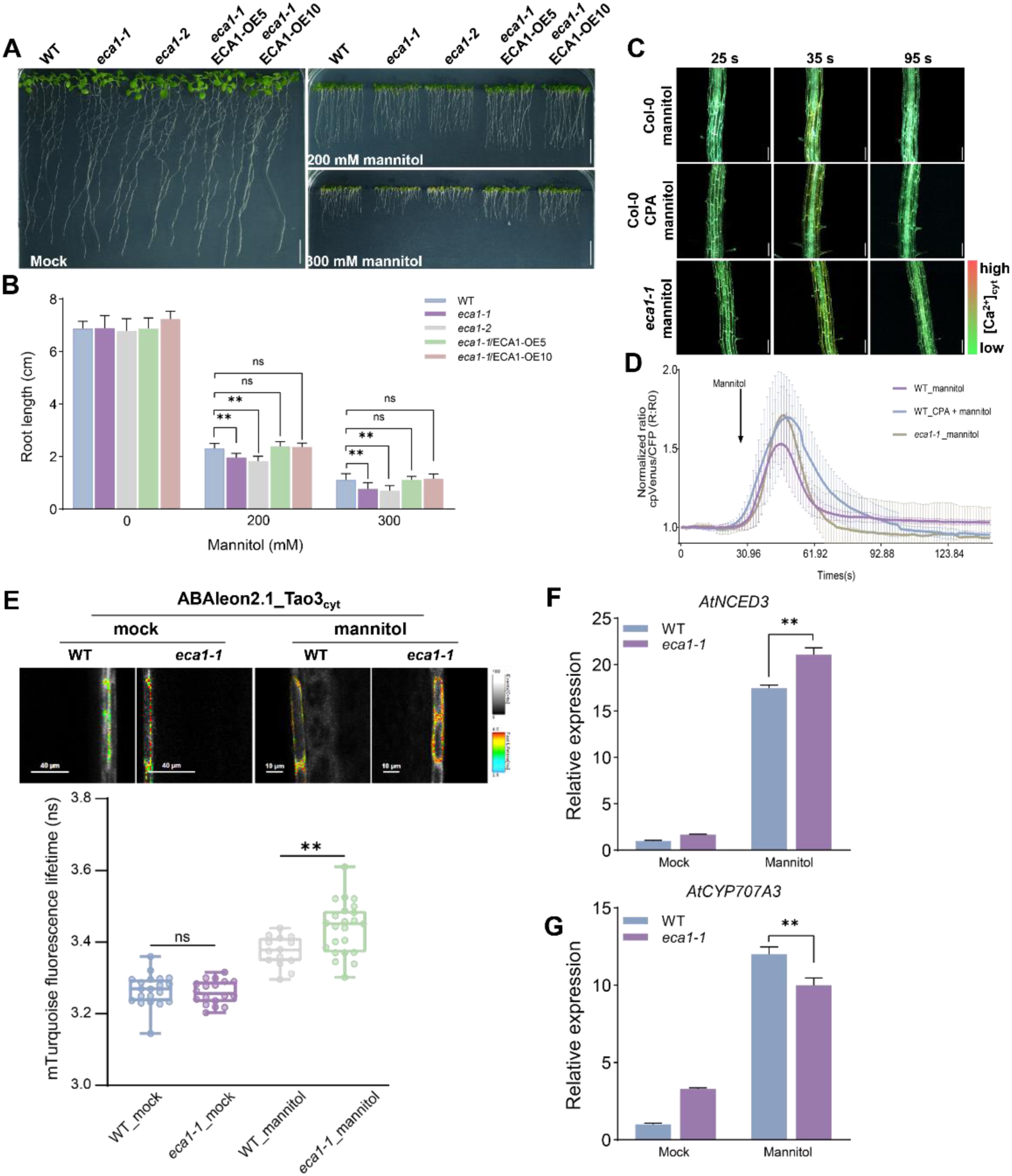
| ECA1 regulates root elongation responses to osmotic stress via changing [Ca^2+^]_cyt_ and [ABA]_cyt_ levels. **A, B** Representative images (**A**) and root length (**B**) of *Arabidopsis* WT, *eca1* mutants, and complementary lines (*eca1-1/ECA1-OE5* and *eca1-1/ECA1-OE10*) treated with 200, 300 mM mannitol. Data are means ± SD, n = 15. Statistics were performed using one-way ANOVA followed by Dunnett’s multiple comparisons test within the same mannitol concentration treatment (\*\**P* < 0.01; ns, not significant). **C, D** Representative images (**C**) and normalized ratio values (ΔR:R0) of [Ca^2+^]_cyt_ changes (**D**) in WT and *eca1-1* sensor lines treated with mannitol (400 mM). WT sensor lines pretreated with CPA were included for comparison. The ratio graphs represent the average ±SD, n = 3. **E** Representation FLIM images and data showing the changes in osmotic stress-enhanced ABA_cyto_ level in *eca1-1* mutant. The ABA sensor expressing lines in WT and *eca1-1* mutant background exposed to mannitol (300 mM) for 4 h. Fluorescence lifetime data are shown as box plot with original data points. Statistics were performed using one-way ANOVA followed by Dunnett’s multiple comparisons test between treatments (\*\**P* < 0.01; ns, not significant). **F, G** Relative expression of *NCED3* (F) and *CYP707A3* (G) in WT and the *eca1-1* mutant upon mannitol treatment. Seedlings were transferred onto ½ MS medium containing 200 mM mannitol for 1 hour before harvested for gene expression analysis. Values are means ± SD, n = 3. Statistics were performed using Students’ *t* test (\*\**P* < 0.01).

To test whether the hypersensitivity to osmotic stress in the *eca1* mutants is related to alterations in [Ca^2+^]cyt and [ABA]cyt levels, we generated calcium and ABA sensor lines in the *eca1-1* mutant background. Notably, the osmotic stress-elicited [Ca^2+^]cyt transients in the *eca1-1* mutant was similarly amplified as those in WT lines co-treated with CPA and mannitol (Fig. 2C, D). Consistently, cytosolic ABA levels in the *eca1-1* mutant were substantially enhanced compared to those in WT sensor lines upon mannitol treatment (Fig. 2E). Further gene expression analysis revealed significantly enhanced and reduced expression of the ABA biosynthesis gene *NCED3* and catabolic gene *CYP707A3*, respectively, in the *eca1-1* mutant triggered upon mannitol (Fig. 2F, G). These results suggest that ECA1 participates in osmotic stress-inhibited root elongation via mediating [Ca^2+^]cyt and [ABA]cyt levels.

### ECA1 interacts with calcium-sensing kinases CPK2/6/11

Regulation of Ca^2+^ channels and pumps by protein kinases represents an important mechanism underlying cellular Ca^2+^ homeostasis^43, 44^. We hypothesized that ECA1 may act as a direct target of calcium sensing protein kinases. Previous *in vitro* phosphorylation analysis of the CPK family showed that the N-terminal Ser19 and Ser22 sites of ACA8 can be phosphorylated by CPK16^45^. This prompted us to focus on the CPK family and screened all the 34 members in the family for potential interactions with ECA1 using bimolecular fluorescence complementation (BiFC) assays. This screening revealed interactions of ECA1 with CPK2, CPK6, CPK11, CPK12, CPK28, and CPK32 (Supplementary Fig. 3). Our phylogenetic analysis using maximum likelihood method divided the CPK family proteins into four groups: group A, B, C, and D (Supplementary Fig. 3). Interestingly, four among the six CPKs that interacted with ECA1 fall into group D (Supplementary Fig. 3), suggesting a binding preference of ECA1 for CPKs in this group.

Considering the stronger interactions of ECA1 with CPK2, CPK6 and CPK11 shown by their stronger fluorescence signals in BiFC assays (Supplementary Fig. 3), we chose these three CPKs for further study. Co-expression of the ER marker CNX-RFP with CPK2/6/11 and ECA1 revealed clear co-localized fluorescent signals in tobacco leaves (Fig. 3A), suggesting the interactions between ECA1 and CPK2/6/11 occurred at the ER membrane/cytosol interface. Co-immunoprecipitation (Co-IP) assays using HA beads showed that ECA1-HA immunoprecipitated CPK2/6/11-GFP (Fig. 3B). Furthermore, fluorescence resonance energy transfer-fluorescence lifetime image (FRET-FLIM) analysis using ECA1-GFP and CPK2/6/11-RFP as FRET pairs revealed evident reductions in GFP fluorescence lifetime in cells co-expressing ECA1-GFP and CPK2/6/11-RFP in comparison with those expressing ECA1-GFP and CPK2/6/11-HA (Fig. 3C). These results show that ECA1 can be targeted by multiple CPKs.

**Fig. 3.**
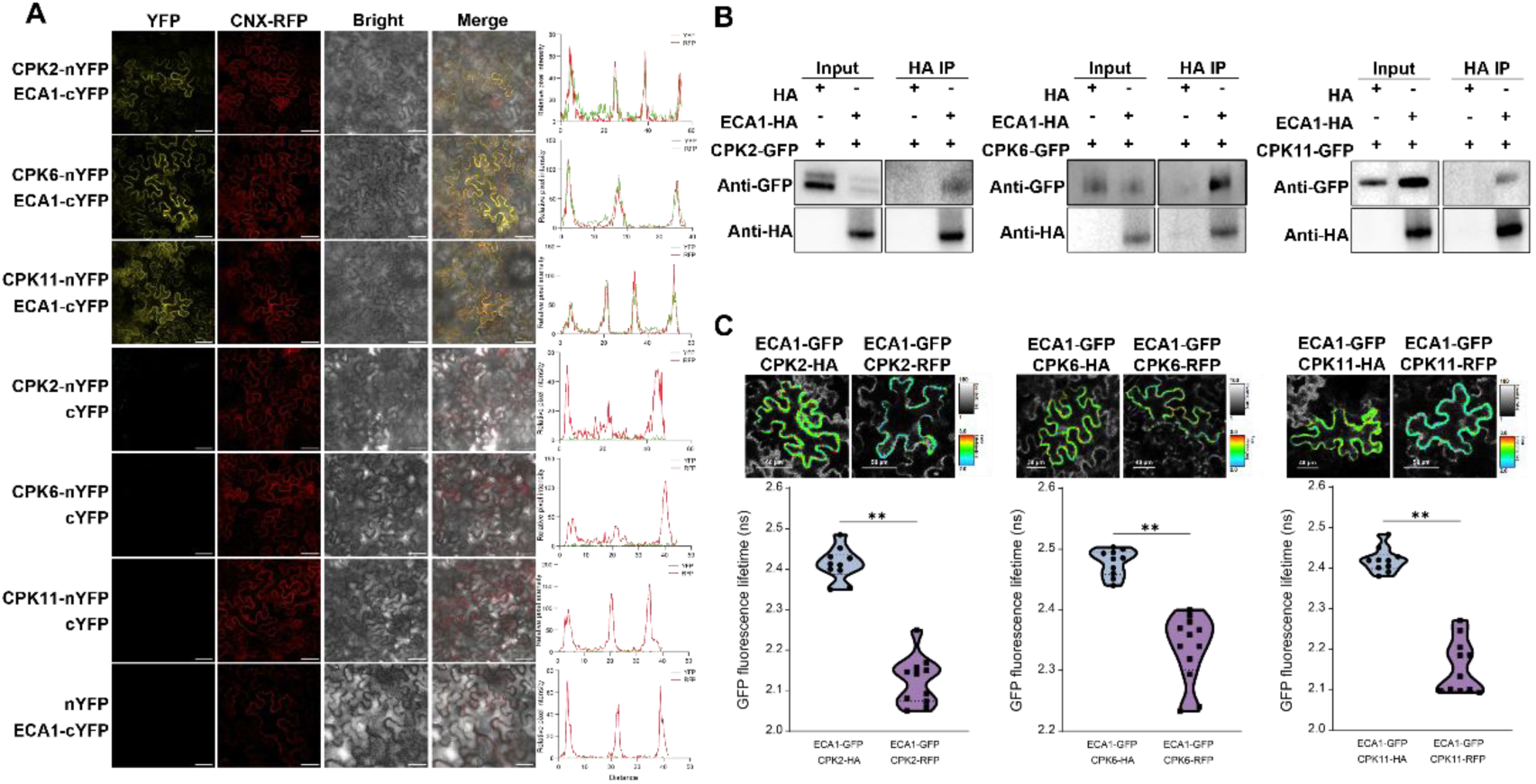
| ECA1 interacts with CPK2/6/11. **A** BiFC assay showing the interaction between CPK2/6/11 and ECA1 at the ER membrane. The ER RFP marker CNX-RFP was co-expressed in all the BiFC constructs combinations. Along the images are the fluorescent intensity plots showing the co-localization of ECA1 and CPK2/6/11 at the ER membrane. **B** Co-immunoprecipitation assay showing the interactions of ECA1 with CPK2/6/11. ECA1-HA and CPK2/6/11-GFP were co-transfected in tobacco leaves and immunoprecipitated with anti-HA beads. Immunoblots were probed with antibodies to detect the presence of ECA1-HA (α-HA) and ECA1-CPK2/6/11 interactions (α-GFP). **C** FRET-FLIM assay revealing the interaction between CPK2/6/11 and ECA1. ECA1-GFP was co-expressed with CPK2/6/11-HA or CPK2/6/11-GFP in tobacco leaves. Data are shown as FLIM images and violin plots. Statistical analysis was performed using Students’ *t* test (\*\**P* < 0.01).

### CPK2/6/11 interact and phosphorylate ECA1 at Ser5 residue

ECA1 resides on the ER membrane with both the N- and C-terminus facing the cytoplasm^46^. To determine with which terminus of ECA1 that the three CPKs are associated, we generated the full-length, N-terminal, and C-terminal fragment of ECA1 fusion proteins and tested their interactions with CPKs both *in vitro* and *in vivo*. Split Luciferase complementation assays (Split-LUC) revealed that the strongest luciferase signals occurred when co-expressing CPK2/6/11-nLuc with ECA1-N-cLuc in comparison to the tabacco detectable luciferase activities in leaf areas expressing CPK2/6/11-nLuc with ECA1-C-cLuc (Fig. 4A-C), indicating that CPK2/6/11 interact with the N-terminal region of ECA1. *In vitro* pull-down assays further corroborated the specific interactions of CPK2/6/11 with the N-terminus ECA1 shown by the fact the purified CPK2/6/11 were only detected when incubated with ECA1-N-GST but not with ECA1-C-GST (Fig. 4D, F).

**Fig. 4.**
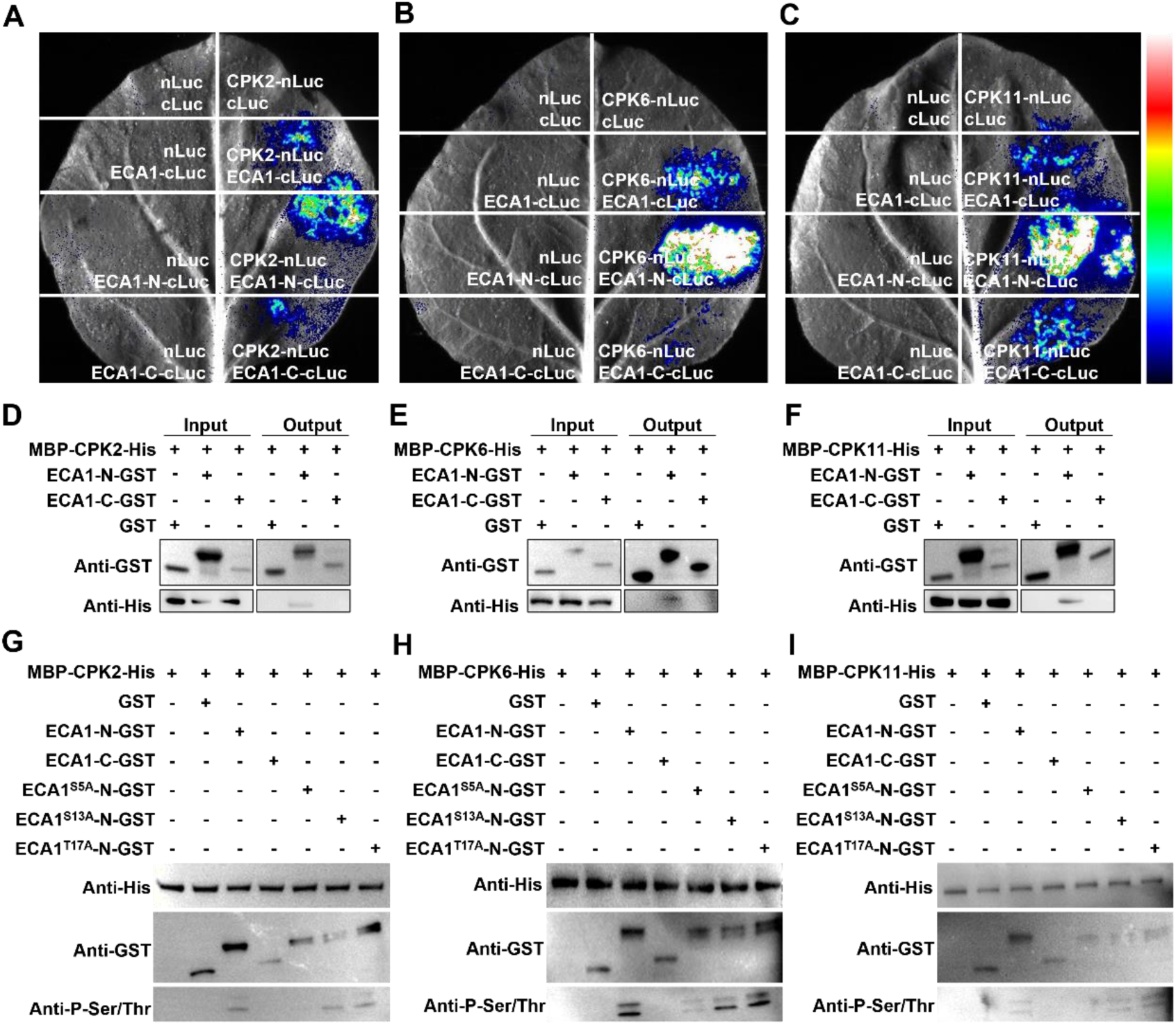
| CPK2/6/11 interact with and phosphorylate the N-terminal of ECA1 at Ser5. **A-C** Split-LUC assays showing the interaction between CPK2 (**A**), CPK6 (**B**), CPK11 (**C**) and ECA1. CPK2/6/11 fused to nLUC (CPK2/6/11-nLUC) were co-transfected with ECA1-N-cLUC or ECA1-C-cLUC in *N. benthamiana* leaves. The empty nLUC was co-expressed with the full-length, N-terminus or C-terminus of ECA1 fused to cLUC as controls. Luciferase images of *N. benthamiana* leaves were taken 48 hours after cultivation. The pseudo-color scale bar shows colors from blue to red, indicating the intensity of the interaction low to high. **D, F** Pull-down assays showing the interactions between CPK2 (**D**), CPK6 (**E**), or CPK11 (**F**) and the N- and C-terminus of ECA1. CPK2/6/11 fused to MBP-His (MBP-CPK2/6/11-His) were purified and incubated with ECA1-N-GST or ECA1-C-GST. CPK2/6/11 and ECA1 were detected using anti-His and anti-GST antibodies. **G-I** *In vitro* phosphorylation assay showing the phosphorylation of the N-terminus of ECA1 at Ser5 by CPK2 (**G**), CPK6 (**H**), or CPK11 (**I**). Phospho-dead variants of ECA1 were generated by mutation of serine/threonine residues at 5, 13 and 17 to alanine (A).

CPKs are typically characterized by their abilities to phosphorylate downstream substrates at serine or threonine residues, thereby regulating plant growth and stress responses^9, 10, 11^. *In vitro* phosphorylation assays confirmed the phosphorylation of ECA1 by CPK2/6/11 at the N-but not the C-terminus (Fig. 4G, H). To further pinpoint the potential phosphorylation sites on ECA1 that are targeted by CPK2/6/11, we first carried out a site query on ECA1 using Plant PTM database (https://www.psb.ugent.be/webtools/ptm-viewer/protein.php?id=AT1G07810.1). It was found that the N-terminal residues Ser5, Ser13, and Thr17 of ECA1 displayed relatively high scores (Supplementary Fig. 4A), indicating these residues as potential phosphorylation sites for CPKs. The three-dimensional structure of the ECA1 protein predicted the spatial locations of these three residues (Supplementary Fig. 4B). Mutation of the serine/threonine residues to Ala resulted in non-phosphorylation of ECA1, hence no CPKs-mediated phosphorylation would occur when incubating the CPK2/6/11-His with the phospho-dead variants of ECA1. This is indeed the case when CPK2/6/11 were incubated with ECA1-N^S5A^ (Fig. 4G-H). Nonetheless, substitution of S13 or T17 with Ala did not alter the phosphorylation at the N terminus of ECA1 (Fig. 4G-H), showing that CPK2/6/11 specifically target ECA1 at the Ser5 residue for phosphorylation modification.

### Phosphorylation of ECA1 at ser5 is vital for its calcium transport activity

Protein phosphorylation and dephosphorylation serve as pivotal molecular switches in plant signaling by altering protein stability or channels/pumps activities^47^. To determine whether ECA1 phosphorylation alters its stability or its activity *in vivo*, we first treated ECA1-GFP seedlings with mannitol and checked protein accumulation in response to treatment. It was found that whilst mannitol treatment did not change the accumulation of the ECA1-GFP protein itself, it triggered evident fluctuations in phosphorylation of ECA1 within 1 h upon treatment (Supplementary Fig. 5). Moreover, we generated a constitutive-phosphorylation variant of ECA1 (ECA1^S5D^-HA) in parallel with the non-phosphorylated variant (ECA1^S5A^-HA). Expressing the variants in *Arabidopsis* protoplasts revealed a slightly enhanced accumulation of ECA1^S5D^-HA under basal but not osmo-stressed condition compared with the wild-type ECA1 and non-phosphorylated ECA1 (Supplementary Fig. 6). These results suggest that phosphorylation of ECA1 has limited effect on its protein stability.

Next, we examined if ECA1 phosphorylation at Ser5 affects its activity for pumping Ca^2+^. Previous heterologous yeast complementation assays using the yeast mutant strain K616, which is known for its incompetence to grow in calcium-depleted medium^48^, have shown that *Arabidopsis* ECA1 can restore the growth of the yeast strain K616 on EGTA^34^. Introduction of both the non-phosphorylated (ECA1^S5A^) and constitutively phosphorylated form (ECA1^S5D^) of ECA1 into the K616 strain showed that yeast cells expressing ECA1^S5D^ grew similarly as the wild-type ECA1 on EGTA-supplemented medium (Fig. 5A, B). Notably, the ECA1^S5A^-expressing cells exhibited a growth rate that resembles the K616 strains without any pump introduction on the EGTA medium (Fig. 5A, B). These results suggest that phosphorylation of ECA1 at ser5 is vital for its channel activity to promote [Ca^2+^]cyt efflux.

**Fig. 5.**
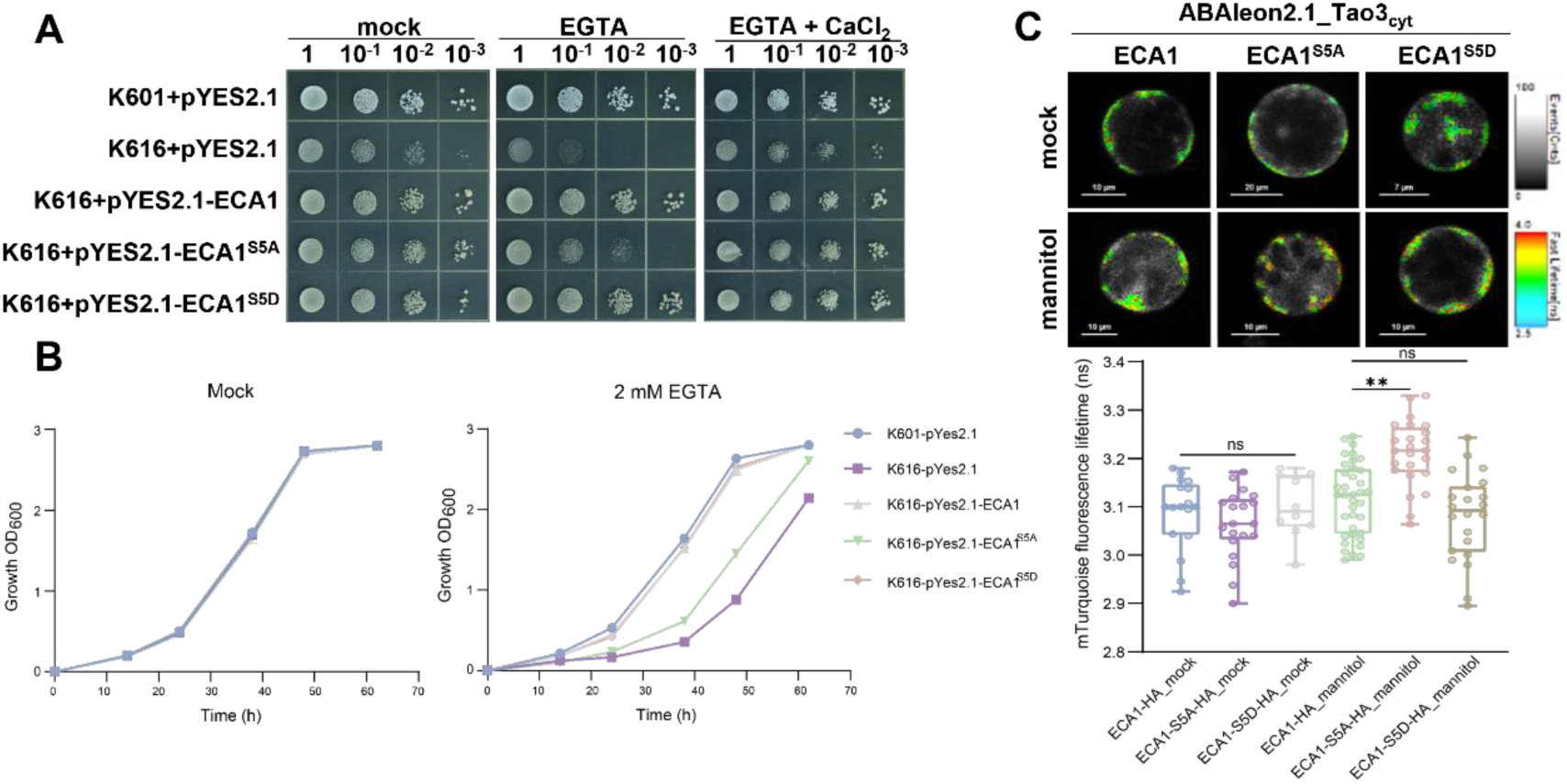
| Phosphorylation of ECA1 at ser5 is vital for its calcium pumping activity. **A** Functional analysis of ECA1 phosphorylation-mediated calcium transport in yeast. The yeast strain K616 was transformed with the empty pYES2.1 vector, pYES2.1 containing wild-type ECA1, ECA1^S5A^, or ECA1^S5D^ and grown on medium (SD/-Ura) supplemented with EGTA (20 mM) or EGTA + CaCl_2_ (10 mM) for 3-5 days. The strain K601 cells transformed with the empty pYES2.1 vector were included as the positive control. From left to right is a 10-fold dilution series of yeast cultures. **B** Quantification of the growth of yeast cells on basal medium (mock) or EGTA supplemented medium at the indicated timepoints. **C** Representation FLIM images and data showing the changes of cytosolic ABA level in Arabidopsis protoplasts exposed to mannitol. The protoplasts co-expressing the ABA sensor ABAleon2.1_Tao3_cyt_ with ECA1-HA, ECA1^S5A^-HA, or ECA1^S5D^-HA were imaged at 2 h upon mannitol treatment (300 mM). Fluorescence lifetime data are shown as box plot with original data points. Statistics were performed using one-way ANOVA followed by Dunnett’s multiple comparisons test between treatments (\*\**P* < 0.01; ns, not significant).

As alterations in [Ca^2+^]cyt positively impacts ABAcyto under osmotic stress (Fig. 1) and ECA1 plays an important role in maintaining osmotic stress-trigged [Ca^2+^]cyt surges (Fig. 2), we suspected that the phosphorylation status of ECA1 may also influence osmotic stress-induced ABAcyto accumulation. Co-expression of cytosolic ABA sensor with wild-type ECA1, ECA1^S5A^, or ECA1^S5D^ in *Arabidopsis* protoplasts revealed that cells expressing ECA1^S5A^ displayed notably higher ABA levels in the cytosol compared to those expressing ECA1 or ECA1^S5D^ upon mannitol treatment (Fig. 5C). These results strongly support that the phosphorylation of ECA1 at ser5 mediates its calcium transport activity, thereby influencing ABAcyto levels under osmotic stress conditions.

### CPK2/6/11 act synergistically with ECA1 in osmotic stress response in *Arabidopsis*

Given that CPK2/6/11 directly target and phosphorylate ECA1 (Fig. 3 and 4), we hypothesized that mutation of CPK2/6/11 would keep ECA1 from the catalytic phosphorylation, thus resulting in compromised regulation of cellular [Ca^2+^]cyt and ABAcyto under osmotic stress conditions. To test this, we crossed *cpk2*, *cpk6*, and *cpk11* mutants with the calcium sensor YC3.6, generating Ca^2+^ sensor lines in the *cpk* mutant background. Treatment of the Ca^2+^ sensor lines with mannitol revealed [Ca^2+^]cyt transients in the *cpk2* and *cpk6* mutant that were similar to that in Col-0, whereas the [Ca^2+^]cyt surged to a notably higher extent in *cpk11* (Fig. 6A). Accordingly, expression of the cytosolic ABA sensor ABAleon2.1_Tao3cyt in protoplasts derived from *cpk* mutants revealed similarly enhanced ABAcyto levels in the wild-type Col-0, *cpk2* and *cpk6* mutants, whereas the *cpk11* mutant accumulated a notably higher amount of ABAcyto upon osmotic treatment (Fig. 6B). These results indicate a redundancy of CPKs and that CPK11 may play a dominant role in mediating cellular Ca^2+^ and ABA homeostasis.

**Fig. 6.**
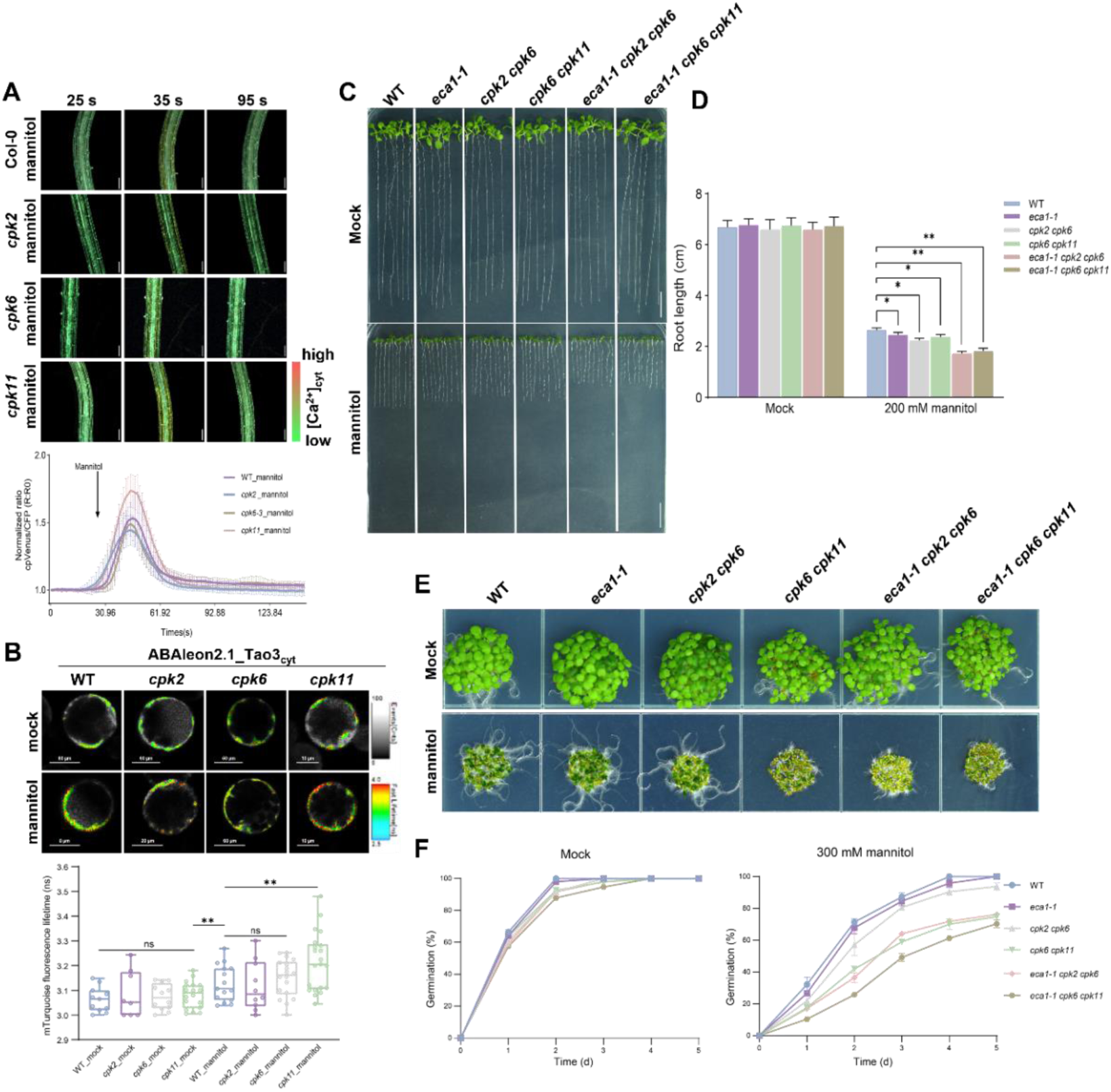
| CPK2/6/11 function with ECA1 in osmotic stress response in *Arabidopsis*. **A** Representative images and normalized ratio values (ΔR:R0) of [Ca^2+^]_cyt_ changes in Col-0, *cpk2*, *cpk6* and *cpk11* mutant expressing the Ca^2+^ sensor YC3.6 treated with mannitol (400 mM). WT sensor lines pretreated with CPA were included for comparison. The arrow indicates the application of mannitol. The ratio graphs represent the average ±SD, n = 3. **B** Representation FLIM images and data showing the changes of cytosolic ABA level in Col-0, *cpk2, cpk6* and *cpk11* protoplasts expressing the ABA sensor ABAleon2.1_Tao3_cyt_ when exposed to 300 mM mannitol. FLIM data are shown as box plot with original data points. Statistics were performed using two-way ANOVA followed by Dunnett’s multiple comparisons test between genotypes within the same treatment (\*\**P* < 0.01; ns, not significant). **C, D** Representative images (**C**) and root length (**D**) of *Arabidopsis* WT Col-0, *eca1-1*, *cpk2 cpk6, cpk6 cpk11*, *eca1-1 cpk2 cpk6* and *eca1-1 cpk6 cpk11* treated with 200 mM mannitol. Data are means ± SD, n = 15. Statistics were performed using one-way ANOVA followed by Dunnett’s multiple comparisons test within the mannitol treatment (\*\**P* < 0.01; \**P* < 0.05). **E, F** Representative images (**E**) and germination (**F**) of *Arabidopsis* WT Col-0, *eca1-1*, *cpk2 cpk6, cpk6 cpk11*, *eca1-1 cpk2 cpk6* and *eca1-1 cpk6 cpk11* treated with 300 mM mannitol.

To determine if CPK2/6/11 act together with ECA1 for controlling stress responses, we crossed *eca1-1*, *cpk2*, *cpk6*, and *cpk11* mutant plants to produce double (*cpk2 cpk6*, *cpk6 cpk11*) and triple mutants (*eca1-1 cpk2 cpk6*, *eca1-1 cpk6 cpk11*) and tested their responses under osmotic stress conditions. Mannitol treatment considerably inhibited root growth, which was exacerbated in *eca1-1* single, *cpk2 cpk6* and *cpk6 cpk11* double mutant (Fig. 6C, D). Moreover, the triple mutants *eca1-1 cpk2 cpk6* and *eca1-1 cpk6 cpk11* was even more sensitive to mannitol treatment than the double mutants, with the shortest roots grown on mannitol-containing medium (Fig. 6C, D). In addition, seed germination assay showed that whilst both the *eca1-1* and *cpk2 cpk6* mutant displayed a similarly impeded germination by mannitol treatment, the double mutant *cpk6 cpk11* and the triple mutants *eca1-1 cpk2 cpk6* and *eca1-1 cpk6 cpk11* showed a notably arrested germination compared to wild-type Col-0 (Fig. 6E, F). These results together support a synergistic effect of CPK2/6/11 and ECA1 in modulating osmotic stress responses.

## Discussion

Ca^2+^ acts as a ubiquitous intracellular signal that controls almost every aspect of cellular functions. Maintaining a balanced Ca^2+^ homeostasis is crucial as excess of [Ca^2+^]cyt triggers cell cytotoxicity^49^. Cytosolic Ca^2+^ dynamics is finely coordinated by an effective Ca^2+^ extrusion system that involves Ca^2+^-ATPases, Ca^2+^/H^+^ exchangers, as well as Ca^2+^-permeable channels^16^. There have been extensive studies on Ca^2+^ influx and efflux across the PM during plant growth and fitness, with few attention being given to Ca^2+^ pumping across organellar membrane and its control of downstream stress responses^50^. This study has unveiled an intracellular signaling cascade that underlies cellular Ca^2+^ and ABA homeostasis for plant resilience to osmotic stress.

Our pharmacological assays using general Ca^2+^ channel blocker LaCl3 revealed precluded cytosolic Ca^2+^ transients without influencing root elongation under osmotic stress (Fig. 1A-D). Given that LaCl3, as a representative rare earth element, elicits multifaceted signaling pathways, the unaltered root growth under LaCl3 and mannitol treatments could be explained by a combined effect of LaCl3 on Ca^2+^, ROS and/or jasmonic acid signaling^51^. In contrast, the ECA1-specific inhibitor CPA notably enhanced plant osmosensitivity, accompanied with augmented and sustained [Ca^2+^]cyt elevation and enriched ABAcyto accumulation (Fig. 1). These point to a fact that whilst reductions in [Ca^2+^]cyt is not necessarily associated with hypo- or insensitivities to stress, improper [Ca^2+^]cyt removal is very likely to cause less tolerant plants. The impaired root growth with magnified Ca^2+^ transients and increased ABAcyto levels in the *eca1* mutants (Fig. 2A-E) further emphasize the fundamental importance of [Ca^2+^]cyt extrusion via ECA1 for plant environmental adaptation.

Ca^2+^ and ABA signaling form tight signaling networks in diverse stress responses^52^. We found that CPKs/ECA1-mediated [Ca^2+^]cyt oscillations are consistently associated with [ABA]cyto alterations (Fig. 1C-E, 2C-E and 6A, B), with Ca^2+^ elevations likely occurring upstream of ABAcyto accumulation given the counteraction of CPA-exacerbated root growth inhibition in ABA synthesis mutants (Fig. 1F). These suggest CPKs/ECA1 contribute to plant osmo-stress tolerance via mediating cellular ABA homeostasis. Studies have shown that Ca^2+^-related proteins are implicated in stress responses by targeting components in ABA signaling^10, 11, 53, 54^. Thus, the pleiotropic effects of Ca^2+^ signaling reflect an intimately controlled ‘Ca^2+^-ABA’ hub for guiding optimal plant growth under adverse conditions.

As the major active extrusion system that ensures cytosolic Ca^2+^ clearance, Ca^2+^-ATPases belonging to PIIB-type are known to be regulated in a built-in autoinhibitory mechanism^23, 55^. We present here that a PIIA-type Ca^2+^-ATPase, ECA1, is directly targeted and phosphorylated by CPK2/6/11 upon osmo-stress sensing, thereby facilitating cytosolic Ca^2+^ removal and restoring [Ca^2+^]cyt to low resting levels (Fig. 3-5). Osmotic stress-triggered root growth inhibition was further exacerbated in the triple mutants *eca1-1 cpk2 cpk6* and *eca1-1 cpk6 cpk11* than that in the single *eca1-1* or double *cpk* mutants (Fig. 6C-F), supporting a collaborative function of ECA1 and CPK2/6/11 in plant osmosensitivity. These findings fill the long-standing knowledge gap between regulation of PIIA-type Ca^2+^-ATPases and plant stress tolerance by showing an intracellular mechanism involving CPK2/6/11-mediated phosphorylation and activation of ECA1 that is directly coupled to plant osmosensitivity via regulating cellular Ca^2+^ and ABA homeostasis. We propose a working model that highlights CPKs/ECA1 as an intracellular calcium sensing machinery for precise control of plant stress tolerance (Fig. 7). Osmotic stress triggers a striking amount of Ca^2+^ influx into the cytoplasm, activating a number of Ca^2+^-sensing proteins like CPKs that are recruited to different intracellular membranes. Three CPKs, CPK2/6/11, are targeted to the ER membrane via interaction with ECA1 and phosphorylate it, which activates the calcium pump activity of ECA1, refraining plants from [Ca^2+^]cyt overload and hypersensitive response to stress. Given that the PM- and ER membrane-associated ACAs and the tonoplast-localized CAXs are also regulated by Ca^2+^-sensing proteins^25, 45, 56^, it is inferred that this Ca^2+^ sensory protein-mediated phosphorylation of calcium channels/pumps at multiple intracellular sites may serve as a broad feedback regulatory mechanism for precise control of cellular Ca^2+^ signature, thereby determining specific physiological outputs. The functional redundancy of ECA1 and CPKs (Fig. 6C-F) in plant osmosensitivity further support an expanded repertoire of Ca^2+^-ATPases that can be targeted by various Ca^2+^-binding protein kinases.

**Fig. 7.**
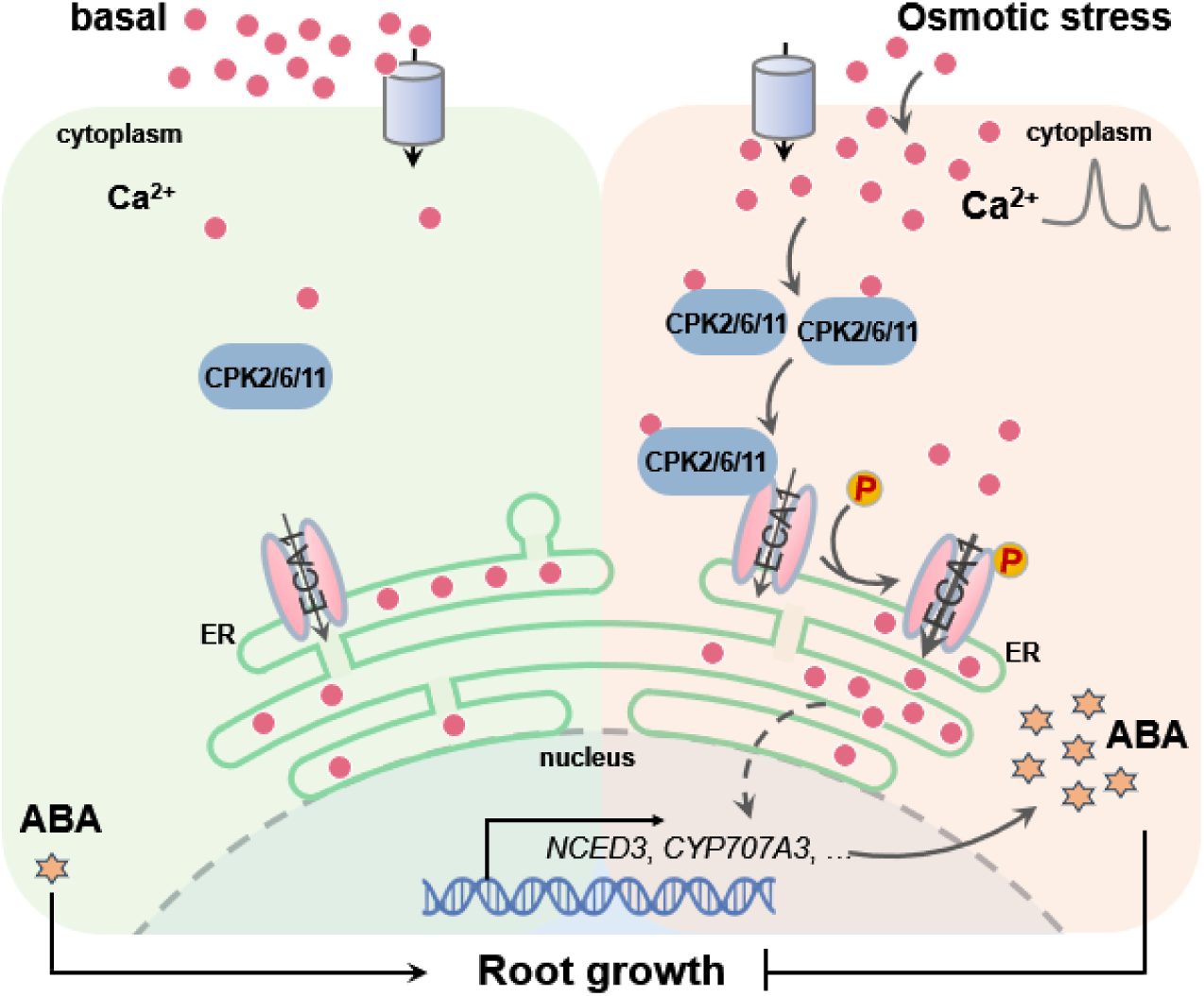
| A working model illustrating CPKs/ECA1-mediated osmotic stress tolerance. Exposure of plants to osmotic stress leads to an increase in cytoplasmic Ca^2+^ concentration, which activates CPK2/6/11. The activated CPK2/6/11 then associate and phosphorylate the ER membrane-localized ECA1 at Ser 5, thereby stimulating ECA1 activity by pumping excessive cytosolic Ca^2+^ into the ER. This restores [Ca^2+^]_cyto_ to a low resting level, ensuring proper plant osmosensitivity.

In ACAs, calmodulin binding sites overlap with the autoinhibitory domain and regulate pumps activities via a bimodular mechanism^23^. It has been shown that Mizu-Kussey 1 (MIZ1) inhibits ECA1 to generate [Ca^2+^]cyt signatures required for root water tracking^35^. In a parallel study, we have shown that CPK2 interacts MIZ1 in a competitive manner with ECA1 (manuscript submitted). Thus, distinct from the autoinhibitory regulation of ACAs, ECAs may be tightly regulated through engaging self-activation and inhibition mechanisms within the Ca^2+^-sensitive triad CPK2-MIZ1-ECA1. How is the triad complex controlled in a temporal-spatial manner with respect to stress response specificity warrant further study.

## Materials and Methods

### Plant materials and growth conditions

*Arabidopsis thaliana* Columbia (Col-0) and *Nicotiana benthamiana*, *Nicotiana tabacum* were used in this study. The T-DNA insertion mutant *eca1-1* (SALK_097881), *cpk2* (SALK_036166) and *cpk11* (SALK_054495) were obtained from the Arabidopsis Biological Resource Center. The *cpk6-3* (SALK_025460) was kindly provided by Dr. Yuanqing Jiang. The *cpk2 cpk6* and *cpk6 cpk11* double mutant were generated by crossing the corresponding single mutant. The *eca1-1 cpk2 cpk6* and *eca1-1 cpk2 cpk6* triple mutant were generated by crossing the corresponding double mutant with *eca1-1*. Seeds were surface sterilized and placed on Petri dishes containing ½-strength Murashige and Skoog (MS) agar (0.8%, w/v) medium and incubated for 2 d at 4°C before transfer to 22°C for germination. To perform osmotic stress treatments, plants were sown on ½ MS medium for 4 days and then transferred to ½ MS with or without 200 mM mannitol for 7 days. Alternatively, seeds were sterilized and directly sown on ½ MS medium with or without 200 mM mannitol, vernalized, and then grown in a light incubator for 7 days. For pharmacological treatments, plants were sown in ½ MS medium for 4 days and then transferred to ½ MS ± 200 mM mannitol, 10 μM CPA (or 50 μM LaCl3) ± 200 mM mannitol for 7 days, respectively. Root length was measured using ImageJ software. To grow plants in the soil, 7-day-old seedlings grown on 1/2 MS medium were transferred to nutrient-rich soil and grown in a greenhouse under long-day conditions (16-hour light/8-hour dark cycle) at 22°C. *N. tabacum* was grown on MS medium supplemented with 2% (w/v) sucrose, 0.5 g L^-1^ MES, and 0.8% (w/v) agar at pH 5.7 in 16/8-h light-dark cycles at 22°C. *N. benthamiana* seeds were planted in nutrient-rich soil and kept in growth chambers under long-day conditions (16-hour light/8-hour dark cycle) at 22°C. After 3 weeks of growth, the plants were used for transformation.

### Construction of Transgenic Plants

For the construction of overexpression vectors, the full-length CDS of ECA1 fragments (3186 bp) was amplified by PCR and cloned into the pGWB505 vector under the control of *CaMV* 35S promoter. *Arabidopsis* transformation with *Agrobacterium tumefaciens* (strain GV3101) was performed by the floral dip method. Homozygous T3 transgenic lines were selected and used for further osmotic stress assay. The calcium indicator YC3.6 (kindly provided by Dr. Alex Costa) was transformed into Arabidopsis Col-0 to generate cytosolic Ca^2+^ sensor lines. Homozygous T3 transgenic lines were selected and further crossed with *eca1-1*, *cpk2*, *cpk6* and *cpk11* mutants for calcium homeostasis analysis.

*eca1-2* mutant was generated using the CRISPR/Cas9 system (Wang et al. 2015). CRISPR guide RNAs (gRNAs) targeting the ECA1 gene were designed using the web tool CRISPR direct (http://crispr.dbcls.jp/)^57^. The CRISPR/Cas9 expression vectors were transformed into the *A. tumefaciens* strain GV3101 and further transformed into Col-0 plants using the floral dip method. Transformants were selected on MS medium supplemented with hygromycin, and their genotypes were confirmed by PCR amplification and sequencing analysis using CRISPR sequencing primers. “Cas9-free” homozygous *eca1-2* mutants were screened in their progeny by culturing them on MS medium supplemented with hygromycin and confirmed by PCR amplification with sequencing analysis.

### Protoplast Isolation and Transfection

Isolation and electrotransfection of tobacco and *Arabidopsis* mesophyll protoplasts were conducted as previously described^40, 58^. Briefly, protoplasts were prepared from perforated tobacco leaves by incubating them overnight in the buffer (3.05 g L^-1^ Gamborg B5 salt medium, 500 mg L^-1^ MES, 750 mg L^-1^ CaCl2.2H2O, 250 mg L^-1^ NH4NO3 adjusted to pH 5.7 with KOH, supplemented with 0.2% w/v macerozyme and 0.4% w/v cellulase) at 25°C in the dark. The protoplasts were then rebuffered through washing three times in 40 mL electrotransfection buffer (137 g L^-1^ sucrose, 2.4 g L^-1^ HEPES, 6 g L^-1^ KCl, 600 mg L^-1^ CaCl2 2H2O adjusted to pH 7.2 with KOH). 500 μL of protoplasts in a total volume of 600 μL electrotransfection buffer were electrotransfected with 2-6 μg plasmid DNA using the Gene Pulser XcellTM (Bio-Rad) with a pulse at 160 V for 10 ms. After transfection, each sample was supplemented with 2 mL incubation buffer and incubated for 16-24 h at 25°C in the dark.

For *Arabidopsis* protoplasts isolation, 2 g of 14-day-old *Arabidopsis* seedlings were sliced into 1-mm strips and placed in 15 mL of filter-sterilized TVL solution (0.3 m sorbitol, 50 mM CaCl2). Next, 20 mL of enzyme solution (0.5 m Sucrose, 10 mM MES-KOH, pH 5.7, 20 mM CaCl2, 40 mM KCl, 1% Cellulase [Onozuka R-10], 1% Macerozyme [R10]) was added, and the mixture was agitated in the dark at room temperature for 16 to 18 hours. The released protoplasts were filtered and pre-washed in W5 solution (0.1% [w/v] GLucose, 0.08% [w/v] KCl, 0.9% [w/v] NaCl, 1.84% [w/v] CaCl2, 2 mM MES-KOH, pH 5.7). The protoplasts, along with any remaining debris, were further filtered and consolidated in a 50-mL centrifuge tube. After centrifuged at 100g for 5 minutes, the protoplasts were washed 2-3 times to remove debris and residual protoplasts. After a 30-minute recovery on ice, the protoplasts were resuspended in MMG solution at the desired concentration. Protoplast aliquots (100 μL) were mixed with plasmid DNA (1 μg in 10 μL) in a 2 mL microcentrifuge tube. Transfection was initiated by adding 110 μL of PEG-calcium solution (40% PEG 4000, 0.2 M mannitol, 100 mM CaCl2). After gentle tapping and a 15-minute incubation at room temperature, the transfection was stopped by diluting the mixture with 440 μL of W5 solution. Transfected protoplasts were collected by centrifugation at 100g for 2 minutes, resuspended in 1 mL of W5 solution, and kept in the dark for the specified time periods before further analysis.

### FRET-Based Ca^2+^ Imaging in *Arabidopsis* Roots

To conduct Ca^2+^ imaging, WT or mutant seeds expressing YC3.6 were germinated and grown in ½ MS medium containing 1% sucrose, 0.6% agar (pH 5.7) for 5 day in a growth chamber under a long-day photoperiod of 16 hours light and 8 hours dark at 22°. Ca^2+^ imaging was performed using a Leica TCS SP8 confocal microscope. Seedlings were mounted on coverslips (40 x 24 mm) using melted gel. Prior to imaging, a small well was created in the gel to allow for the application of solutions, exposing the mature zone of the root. Seedlings were then incubated with control buffer (composed of ½ MS medium, pH 5.8) for 30 minutes before imaging. For mannitol treatment, measurement was taken after using pipettes to replace the buffer solution containing 400 mM mannitol. The YC3.6 Ca^2+^ sensor was excited with the 458-nm line of the argon laser. CFP and cpVenus emissions were detected at 473-505 nm and 526-536 nm, respectively. Image acquisition settings were as follows: scanning speed (400 Hz), image dimension (512 x 512 pixels), pinhole (1 airy unit) and line average (1). Bright-field images were acquired simultaneously using the transmission detector of the microscope. For time-lapse analysis, YFP and CFP images were collected every 1.293 seconds. For quantification, ratio values were normalized to the baseline (R_0_) to obtain relative ratio changes for all measured timepoints (ΔR/R_0_). This normalization was performed using the following equation: ΔR/R_0_ = (R - R_0_)/R_0_.

### Fluorescence lifetime Imaging Microscopy

FLIM recordings of ABA sensor ABAleon2.1_Tao3s were conducted using a Nikon A1R confocal microscope equipped with a PicoHarp time-correlated single-photon counting (TCSPC) module and a PDL800-D multichannel picosecond pulsed diode laser driver (PicoQuant, Berlin, Germany). For the ABAleon2.1_Tao3s sensor, the donor <u>τmT</u> was excited with a 440 nm laser at a 20 MHz pulse frequency. Emissions were recorded at 482/35 nm by TCSPC until a count of at least 200 photons was reached in the brightest pixel, with a total count of at least 10,000 photons. The FLIM imaging method followed the approach described by Zhou (2021). When using GFP vs. RFP as the FRET pair, the donor GFP was excited with a 485 nm laser at 20 MHz pulse frequency. Emission was recorded at 510/35 nm by TCSPC until a count of at least 50 photons was reached in the brightest pixel. At least 10 cells per sample per treatment were recorded. The FLIM data were analyzed using SymphoTime64 v2.0 (PicoQuant, Berlin, Germany) and the calculation method described by Zhou et al. (2021) and Wang et al. (2022).

### Bimolecular Fluorescence Complementation (BiFC)

For BiFC experiments, the CDS of CPK2, CPK6 and CPK11 were amplified by PCR and cloned into pZY101-nYFP, and the CDS of ECA1 was amplified by PCR pZY101-cYFP vectors, respectively, resulting in the generation of CPK2/6/11-nYFP and ECA1-cYFP constructs. The corresponding vector were then transformed into *Agrobacterium tumefaciens* GV3101 cells and injected into *Nicotiana benthamiana* leaves. The YFP fluorescence signal was detected by using a confocal laser scanning microscope (Leica SP8) after 2-day infiltration. The YFP fluorescence was excited with a 514 nm laser, and emission spectra were collected within the range of 530 to 550 nm to detect the YFP signal.

### Split-LUC Complementation Assay

For the Luciferase complementation imaging assays, the full-length CDS of CPK2, CPK6 and CPK11 were amplified by PCR and cloned into pCambia1300-nLUC to obtain CPK2/6/11-nLUC. The full-length CDS of ECA1 and ECA1-N, ECA1-C were amplified by PCR and cloned into pCambia1300-cLUC to obtain ECA1-cLUC and ECA1-N-cLUC, ECA1-C-cLUC. The corresponding vector were then transformed into Agrobacterium tumefaciens GV3101 cells and injected into *Nicotiana benthamiana* leaves. After 2-day incubation period, spray 1 mM Luciferin on the surface of the leaves and then put in dark for 5-10 min The luminescence signals were subsequently detected by CCD imaging system (Tanon, 6100C).

### Co-immunoprecipitation Assay (Co-IP)

For detection the in vivo interaction of CPK2/6/11 and ECA1, the CDS of CPK2, CPK6 and CPK11 were cloned into pGWB505 vector to obtain 35S: CPK2/6/11-GFP. The CDS of ECA1 was cloned into the Pgwb414 vector to obtain 35S: ECA1-HA. CPK2/6/11-GFP and ECA1-HA were transformed into Agrobacterium tumefaciens GV3101 cells. For co-IP in *Nicotiana benthamiana*, leaves from three-week-old soil-grown plants were hand-infiltrated with different pairs of Agrobacterium tumefaciens carrying indicated vectors. Leaf samples were harvested after two days inoculation and liquid nitrogen grinding, subjected to homogenization with IP buffer (50 mM HEPES, PH 7.5, 150 mM KCl, 1 mM DTT, 0.4% Trixton-X 100, 10% Glycerol, protease inhibitor cocktail). Ice bath extraction for 1 hour, then centrifuged at 12,000g for 10 min at 4°C. The supernatant was collected and incubated with anti-GFP mAb-magnetic beads (MBL) gently shaking on a rocker overnight at 4°C. The beads were collected and washed three times with washing buffer (50 mM HEPES, PH 7.5, 150 mM KCl, 1 mM DTT, 0.4% Trixton-X 100). The immunoprecipitated proteins and input proteins were analyzed by immunoblotting using the indicated antibodies. Both anti-GFP (MBL) and anti-HA (Roche) antibodies were used at 1:5000 dilutions.

### Protein Expression and Purification

Plasmids of recombinant proteins were transformed to *Escherichia coli* BL21 (DE3) strain. A single clone of each transformant was selected and incubated in 5 ml of LB liquid medium supplemented with corresponding antibiotics at 37°C for 12 hours and then transferred to 100 ml of LB liquid medium with corresponding antibiotics at 37°C for 2 to 4 hours until the OD 600 reached to 0.6. Unless otherwise indicated, recombinant proteins were expressed in *E. coli* with 0.5 mM isopropyl-d-thiogalactopyranoside (IPTG) for 12 hours at 16°C. GST-tagged recombinant proteins were purified using GST mag-beads (BBI, catalog no. C650031) according to manufacturer’s instructions. His-tagged proteins were purified using His mag-beads (BBI, catalog no. C650033) according to manufacturer’s instructions.

### Pull-down Assay

For GST pull-down assay, CPK2, CPK6 and CPK11 fragments fused with MBP-His. ECA1-N and ECA1-C fragments fused with GST. In brief, 2 μg of MBP-CPK2/6/11-His proteins were incubated with 1.5 μg of ECA1-N/C-GST and immunoprecipitated by GST beads at 4°C for 1 hours. Collect the beads with a magnetic stand. The GST beads was washed four times with PBS buffer and eluted with 10 mM reduced gluthatione. The proteins were separated by SDS-PAGE and immunoblotted with anti-His (MBL), anti-GST (Beyotime Biotechnology).

### *In Vitro* Phosphorylation Assay

For *in vitro* phosphorylation assay, ECA1-N, ECA1-N^S5A^, ECA1-N^S13A^, ECA1-N^T17A^, ECA1-C were amplified and cloned into the pGEX6p-2, CPK2, CPK6 and CPK11 CDSs were amplified and cloned into the pMAL-MBP-His vectors to obtain ECA1-N-GST, ECA1-N^S5A^-GST, ECA1-N^S13A^-GST, ECA1-N^T17A^-GST, ECA1-C-GST, MBP-CPK6-His. In brief, 1.5 μg of MBP-CPK2/6/11-His proteins was incubated separately with 1.5 μg of ECA1-N-GST, ECA1-NS5A-GST, ECA1-NS13A-GST, ECA1-NT17A-GST, ECA1-C-GST in phosphorylation buffer (20 mM Tris-HCl (PH7.5), 5 mM MgCl2, 1 mM DTT, 1 mM CaCl2, 50 μM ATP, protease inhibitor cocktail, Protein phosphatase inhibitors), 30°, 30min. The reaction was terminated by adding SDS loading buffer and incubating at 100° for 5 min. The reaction mixture was analyzed by immunoblotting with indicated antibodies. Anti-phosphoserine/threonine (ECM Biosciences) antibody was used at 1:1000 dilutions.

### Yeast transformation, complementation and growth

The yeast strain K616 was transformed with plasmids pYES2.1-ECA1, pYES2.1-ECA1^S5A^, pYES2.1-ECA1^S5D^, or the empty pYES2.1 vector. As a positive control, the yeast strain K601 was transformed with the empty pYES2.1 vector. The transformed yeast cells were selected on SD/-Ura medium for 3 to 5 days. For complementation studies, single colonies were grown in SD-URA liquid medium, harvested, washed three times with 10 mM EGTA and diluted to an OD600 of 1.0. Ten μL of serial 10-fold dilutions were spotted onto SD-URA plates supplemented with 2% (w/v) galactose, 1% (w/v) raffinose and 20 mM EGTA. The plates were then incubated at 30°C for 3–5 days to assess growth and complementation. For the growth rate experiment, the yeast cells carrying different vectors were grown in liquid medium SD-U or SD-U supplemented with 2 mM EGTA for indicated time.

### RNA Isolation and RT-qPCR Analysis

Total RNA was extracted from plants with ReliaPrep™ RNA Miniprep System (Promega) following the manufacturer’s protocols. For RT-qPCR analyses, total RNA was used for cDNA synthesis by NovoScript^®^Plus All-in-one 1^st^ Strand cDNA Synthesis SuperMix (Novoprotein). The quantitative PCR analysis was performed using an Applied Biosystems 7500 real-time PCR system. The SYBR Green Master Mix (YEASEN) was used for reaction according to the manufacturer’s instruction. Arabidopsis *Actin2* was used as an internal control. The primer sequences for RT-qPCR are listed in Supplemental Table 1.

### Quantification and Statistical Analysis

Data for quantification analyses are presented as mean ± standard error (SE) or standard deviation (SD). Statistical analyses were performed using Students’ *t* test or ANOVA (one-way or two-way) to calculate p values using Prism software (GraphPad Software, La Jolla, CA). Data shown are representative from at least three independent experiments.

## Author contributions

X.L., Y.Z. and J. L. conceived and designed the experiments. X.L. performed the experiments. X.L., Y. Z., W.X. and J. L. analyzed the data and wrote the manuscript. All authors reviewed and approved the manuscript.

## Funding and Acknowledgements

This work was supported by grants from the National Natural Science Foundation of China (32070292 to J.L.), Shenzhen Science and Technology Program (grant no.: KQTD20190929173906742 and ZDSYS20230626091659010 to J.L.). The authors acknowledge the assistance of SUSTech Core Research Facilities and SUSTech IPF Core Facilities.

## Supplementary Information

**Supplementary Fig. 1.**
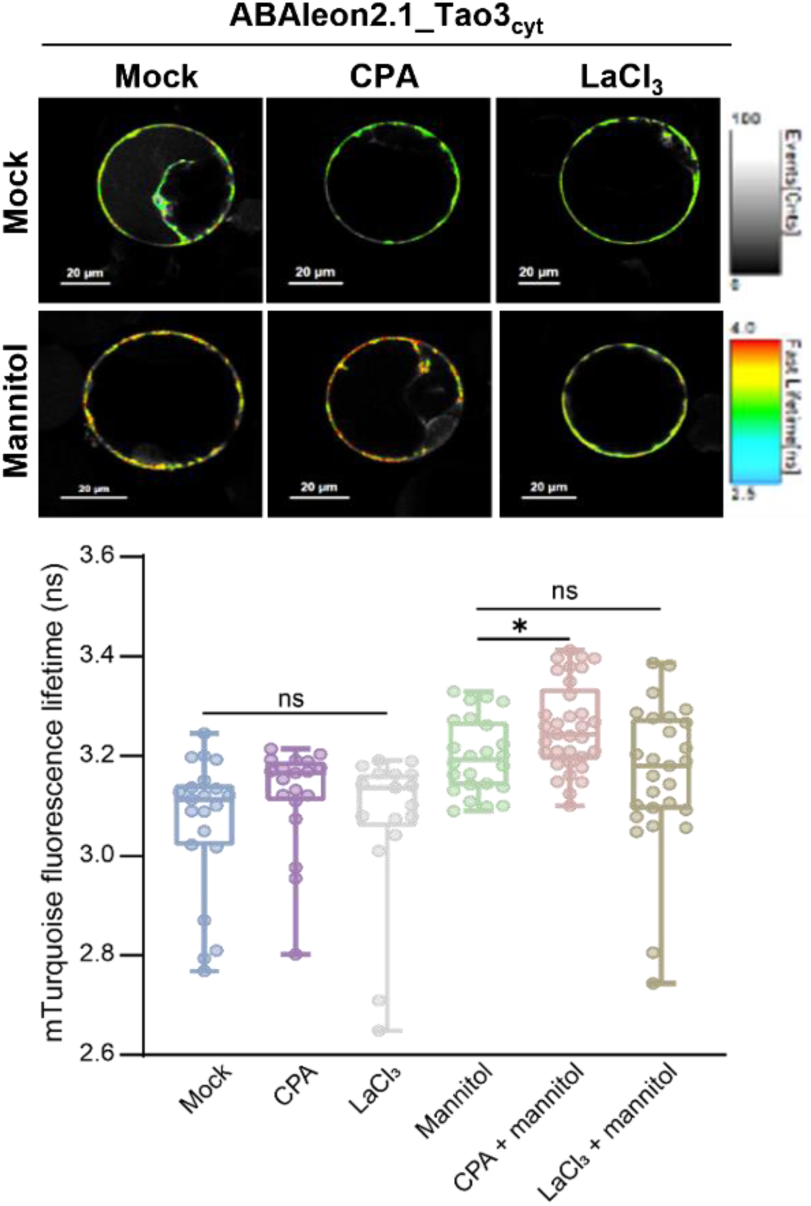
| Effect of Ca^2+^ channel inhibitors on osmotic stress-induced cytosolic ABA accumulation in tobacco protoplasts. Representation FLIM images and data showing the changes of cytosolic ABA level in response to mannitol in the presence or absence of calcium inhibitors. Protoplasts derived from tobacco leaf mesophyll cells expressing the cytosolic ABA sensor ABAleon2.1_Tao3cyt were pretreated with LaCl3 (50 μM) or CPA (5 μM) before exposed to mannitol (300 mM) for 4 h. Fluorescence lifetime data are shown as box plot with original data points. Statistics were performed using one-way ANOVA followed by Dunnett’s multiple comparisons test between treatments (\**P* < 0.05; ns, not significant).

**Supplementary Fig. 2.**
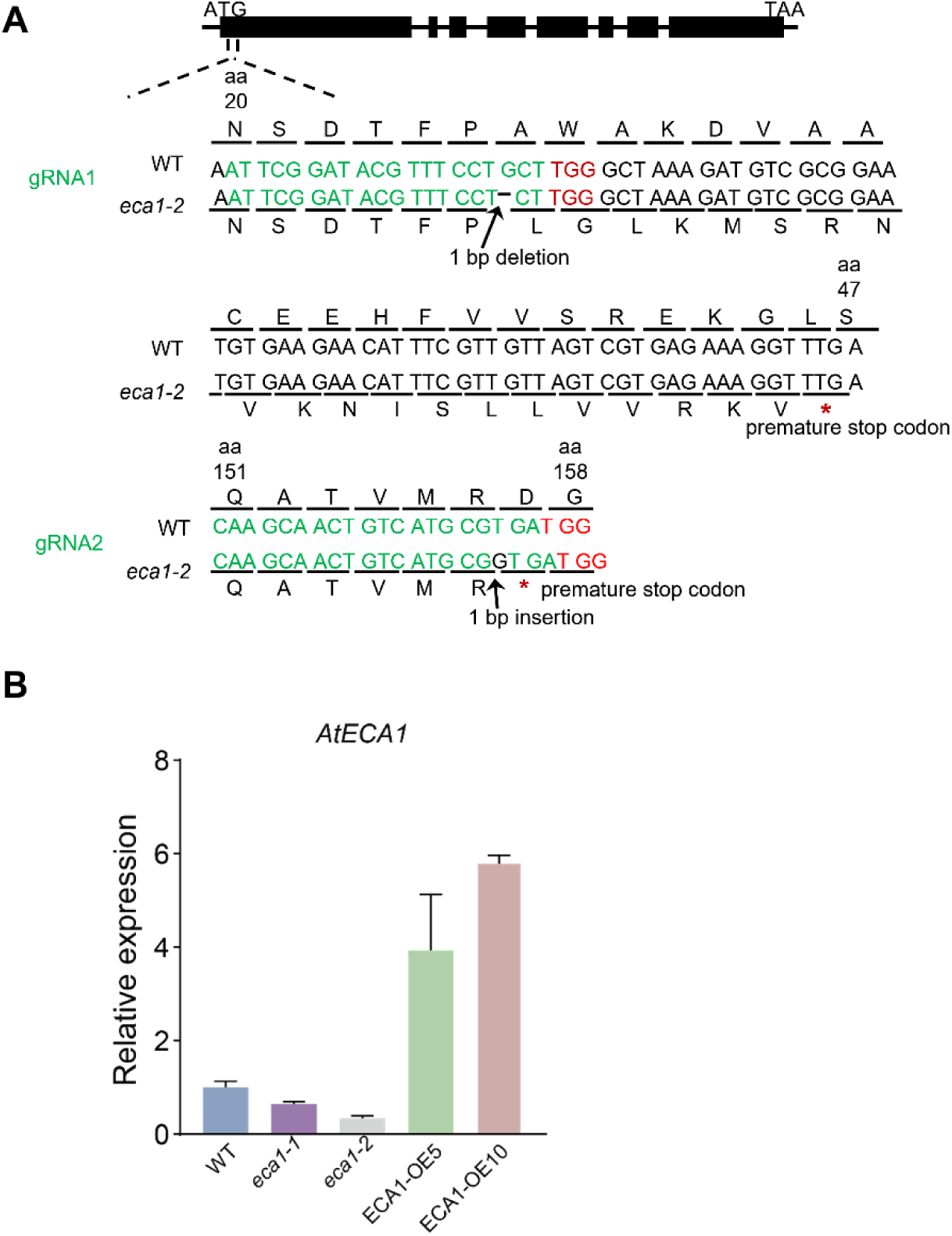
| Generation of the *eca1-2* mutant by CRISPR/Cas9. **A** Targeted areas in the ECA1 gene. Black boxes in the gene body indicate the exon regions, and horizontal lines indicate the intron regions. The *eca1-2* mutant carries a 1-bp deletion at nucleotide position 76, resulting in frameshift mutation and premature stop codon. Green capital letters represent the targeted sequences in ECA1. Red capital letters represent protospacer adjacent motif (PAM) sequences. **B** Quantification of the gene expression level of *ECA1* in *Arabidopsis* Col-0 (WT), *eca1-1*, *eca1-2* and *ECA1* overexpression line (ECA1-OE5, ECA1-OE10). Total RNA extracted from eight-day-old seedlings. Values are means ± SD. n = 3.

**Supplementary Fig. 3.**
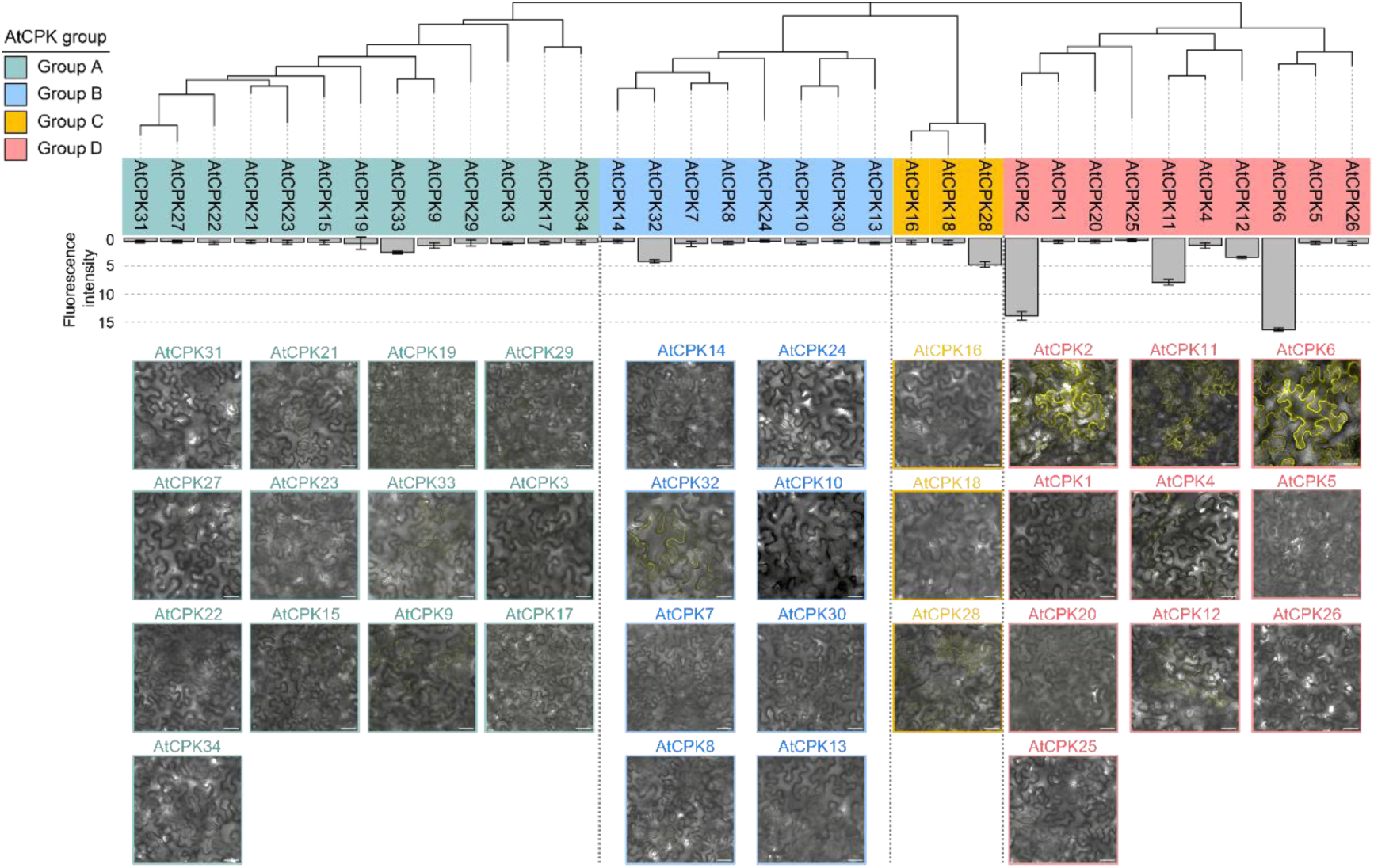
| Screening of ECA1 interaction partners in CPKs family in *N. benthamiana* leaves. The Upper panel shows the phylogenetic tree of CPK proteins that is constructed using the Maximum likelihood (ML) method in MEGA X software. Amino acid sequences of CPKs proteins were downloaded from the NCBI database for this analysis. Four groups in the phylogenetic tree are represented by different colors: A (green), B (blue), C (yellow), and D (red). The middle and lower panel show the YFP fluorescence intensities and interactions between CPKs and ECA1 in BiFC assays, respectively. CPK proteins fused to nYFP (CPK-nYFP) were co-expressed with ECA1 that was fused to cYFP (ECA1-cYFP) and YFP fluorescence was detected under confocal microscopy.

**Supplementary Fig. 4.**
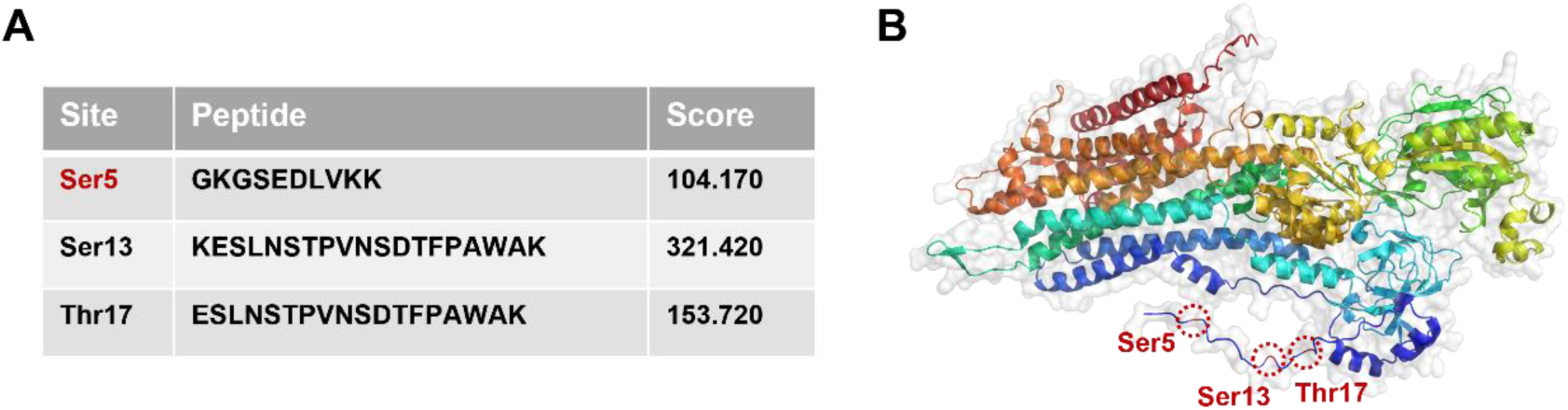
| Prediction of phosphorylation sites of ECA1. **A** Predicted scores of the potential phosphorylation sites at Ser5, Ser13, and Thr17 in ECA1. The query was performed on the Plant PTM database (https://www.psb.ugent.be/webtools/ptm-viewer/protein.php?id=AT1G07810.1) utilizing the ECA1 protein sequence. **B** Protein structure image of ECA1 highlighting potential phosphorylation sites at the cytoplasm-ER interface. The phosphorylation sites Ser5, Ser13, and Thr17 are shown in red. The protein model was generated using PyMOL 2.5.5.

**Supplementary Fig. 5.**
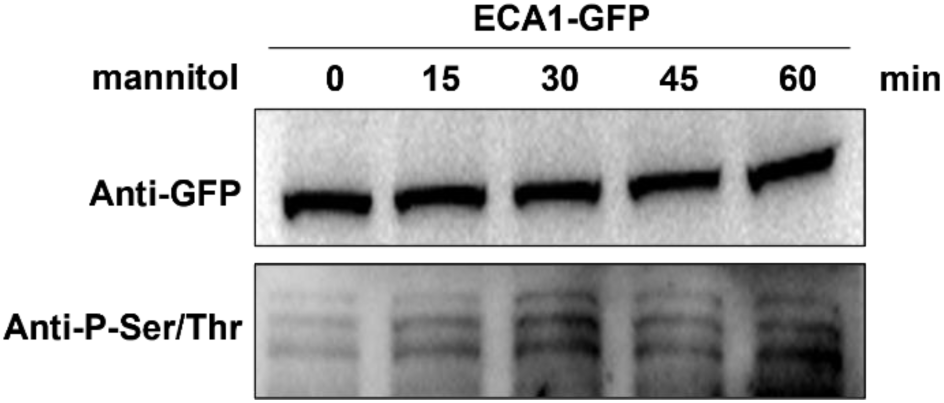
| Phosphorylation of ECA1 in response to osmotic stress. Changes in the phosphorylation level of ECA1 protein under osmotic stress. Seven-day-old transgenic plants overexpressing ECA1-GFP were treated with mannitol (200 mM) for the indicated time periods. The ECA1 protein accumulation and phosphorylation level were detected using anti-GFP and anti-P-Ser/Thr antibodies, respectively.

**Supplementary Fig. 6.**
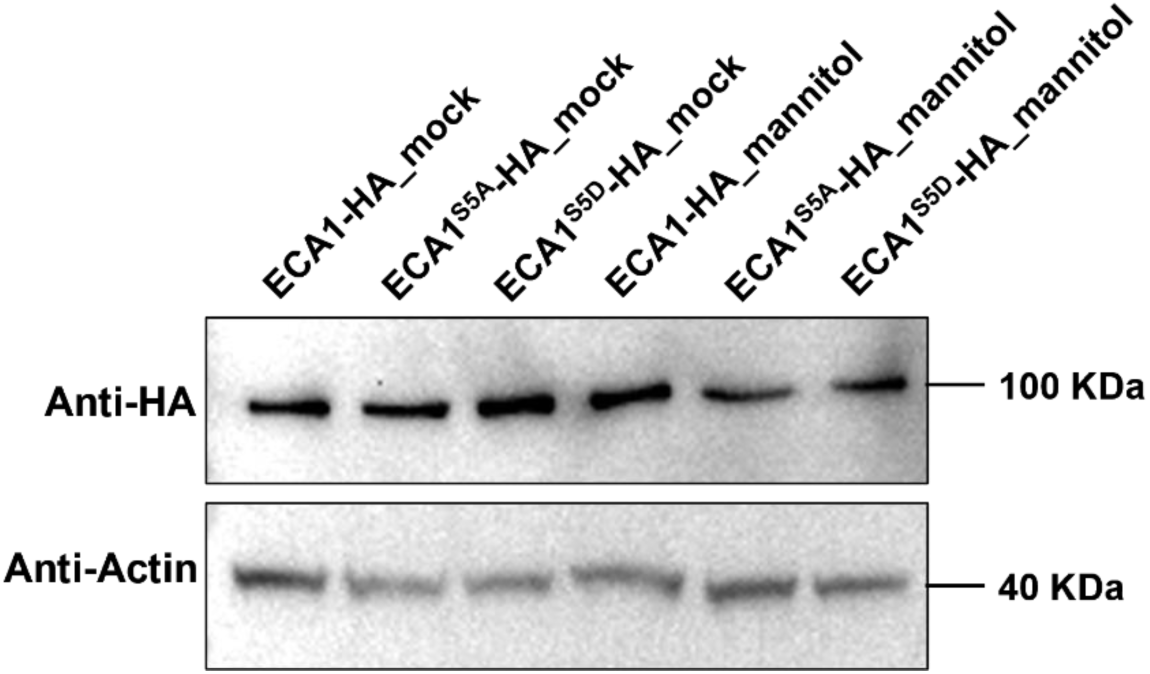
| Effect of ECA1 phosphorylation on its protein stability. The ser5 residue in ECA1 was mutated to alanine (A) and aspartic acid (D) to generate non-phosphorylated and constitutive phosphorylation form of ECA1, respectively. Arabidopsis Col-0 protoplasts were transfected with HA-tagged ECA1 variants and exposed to mannitol (300 mM) for 3 h before harvest for immunoblot analysis. Anti-HA antibody was used to detect the ECA1 proteins. Anti-Actin antibody was included as a loading control.

## References

1. Pei S, et al. Osmosensor-mediated control of Ca^2+^ spiking in pollen germination. Nature 629, 1118–1125 (2024).

2. Schiøtt M, Romanowsky SM, Bækgaard L, Jakobsen MK, Palmgren MG, Harpe JF. A plant plasma membrane Ca^2+^ pump is required for normal pollen tube growth and fertilization. Proc Natl Acad Sci USA 101, 9502–9507 (2004).

3. Leitão N, Dangeville P, Carter R, Charpentier M. Nuclear calcium signatures are associated with root development. Nat Commun 10, 4865 (2019).

4. Stephan AB, Kunz H-H, Yang E, Schroeder JI. Rapid hyperosmotic-induced Ca^2+^ responses in *Arabidopsis thaliana* exhibit sensory potentiation and involvement of plastidial KEA transporters. Proc Natl Acad Sci USA 113, E5242–E5249 (2016).

5. Tian W, et al. A calmodulin-gated calcium channel links pathogen patterns to plant immunity. Nature 572, 131–135 (2019).

6. Yuan F, et al. OSCA1 mediates osmotic-stress-evoked Ca^2+^ increases vital for osmosensing in *Arabidopsis*. Nature 514, 367–371 (2014).

7. Boudsocq M, Sheen J. CDPKs in immune and stress signaling. Trends in Plant Sci 18, 30–40 (2013).

8. Zeng H, Xu L, Singh A, Wang H, Du L, Poovaiah BW. Involvement of calmodulin and calmodulin-like proteins in plant responses to abiotic stresses. Front Plant Sci 6, (2015).

9. Yip Delormel T, Boudsocq M. Properties and functions of calcium-dependent protein kinases and their relatives in Arabidopsis thaliana. New Phytol 224, 585–604 (2019).

10. Zhang H, et al. Arabidopsis CPK6 positively regulates ABA signaling and drought tolerance through phosphorylating ABA-responsive element-binding factors. J Exp Bot 71, 188–203 (2020).

11. Zhu S-Y, et al. Two Calcium-Dependent Protein Kinases, CPK4 and CPK11, Regulate Abscisic Acid Signal Transduction in *Arabidopsis*. Plant Cell 19, 3019–3036 (2007).

12. Gao Q-F, et al. Cyclic nucleotide-gated channel 18 is an essential Ca^2+^ channel in pollen tube tips for pollen tube guidance to ovules in *Arabidopsis*. Proc Natl Acad Sci USA 113, 3096–3101 (2016).

13. Hilleary R, Paez-Valencia J, Vens CS, Toyota M, Palmgren M, Gilroy S. Tonoplast-localized Ca^2+^ pumps regulate Ca^2+^ signals during pattern-triggered immunity in *Arabidopsis thaliana*. Proc Natl Acad Sci USA 117, 18849–18857 (2020).

14. Thor K, et al. The calcium-permeable channel OSCA1.3 regulates plant stomatal immunity. Nature 585, 569–573 (2020).

15. Jaślan D, et al. Voltage-dependent gating of SV channel TPC1 confers vacuole excitability. Nat Commun 10, (2019).

16. Demidchik V, Shabala S, Isayenkov S, Cuin TA, Pottosin I. Calcium transport across plant membranes: mechanisms and functions. New Phytol 220, 49–69 (2018).

17. Kintzer AF, Stroud RM. Structure, inhibition and regulation of two-pore channel TPC1 from Arabidopsis thaliana. Nature 531, 258–264 (2016).

18. Charpentier M, et al. Nuclear-localized cyclic nucleotide-gated channels mediate symbiotic calcium oscillations. Science 352, 1102–1105 (2016).

19. Laohavisit A, et al. Salinity-Induced Calcium Signaling and Root Adaptation in Arabidopsis Require the Calcium Regulatory Protein Annexin1. Plant Physiol 163, 253–262 (2013).

20. Michard E, et al. Glutamate Receptor–Like Genes Form Ca^2+^ Channels in Pollen Tubes and Are Regulated by Pistil d-Serine. Science 332, 434–437 (2011).

21. Ortiz-Ramírez C, et al. GLUTAMATE RECEPTOR-LIKE channels are essential for chemotaxis and reproduction in mosses. Nature 549, 91–95 (2017).

22. Nakagawa Y, et al. Arabidopsis plasma membrane protein crucial for Ca^2+^ influx and touch sensing in roots. Proc Natl Acad Sci USA 104, 3639–3644 (2007).

23. Tidow H, et al. A bimodular mechanism of calcium control in eukaryotes. Nature 491, 468–472 (2012).

24. Huda KMK, Banu MSA, Tuteja R, Tuteja N. Global calcium transducer P-type Ca^2+^-ATPases open new avenues for agriculture by regulating stress signalling. J Exp Bot 64, 3099–3109 (2013).

25. Wang C, et al. Mechanisms of calcium homeostasis orchestrate plant growth and immunity. Nature 627, 382–388 (2024).

26. Sze H, Liang F, Hwang I. DIVERSITY AND REGULATION OF PLANT Ca^2+^ PUMPS: Insights from Expression in Yeast. Annu Rev Plant Physiol Plant Mol Biol 51, 433–462 (2000).

27. Iwano M, et al. A Pollen Coat–Inducible Autoinhibited Ca^2+^-ATPase Expressed in Stigmatic Papilla Cells Is Required for Compatible Pollination in the Brassicaceae. Plant Cell 26, 636–649 (2014).

28. An Y, et al. 5-Aminolevulinic Acid Thins Pear Fruits by Inhibiting Pollen Tube Growth via Ca^2+^-ATPase-Mediated Ca^2+^ Efflux. Front Plant Sci 7, 121 (2016).

29. Zhang J, Zhang X, Wang R, Li W. The plasma membrane-localised Ca2+-ATPase ACA8 plays a role in sucrose signalling involved in early seedling development in Arabidopsis. Plant Cell Rep 33, 755–766 (2014).

30. Zhu X, Caplan J, Mamillapalli P, Czymmek K, Dinesh-Kumar SP. Function of endoplasmic reticulum calcium ATPase in innate immunity-mediated programmed cell death. EMBO J 29, 1007–1018 (2010).

31. Boursiac Y, et al. Disruption of the Vacuolar Calcium-ATPases in Arabidopsis Results in the Activation of a Salicylic Acid-Dependent Programmed Cell Death Pathway. Plant Physiol 154, 1158–1171 (2010).

32. Frei dit Frey N, et al. Plasma Membrane Calcium ATPases Are Important Components of Receptor-Mediated Signaling in Plant Immune Responses and Development. Plant Physiol 159, 798–809 (2012).

33. Wu Z, et al. An Endoplasmic Reticulum-Bound Ca^2+^/Mn^2+^ Pump, ECA1, Supports Plant Growth and Confers Tolerance to Mn^2+^ Stress. Plant Physiol 130, 128–137 (2002).

34. Liang F, Cunningham KW, Harper JF, Sze H. ECA1 complements yeast mutants defective in Ca^2+^ pumps and encodes an endoplasmic reticulum-type Ca^2+^-ATPase in *Arabidopsis thaliana*. Proc Natl Acad Sci USA 94, 8579–8584 (1997).

35. Shkolnik D, Nuriel R, Bonza MC, Costa A, Fromm H. MIZ1 regulates ECA1 to generate a slow, long-distance phloem-transmitted Ca^2+^signal essential for root water tracking inArabidopsis. Proc Natl Acad Sci USA 115, 8031–8036 (2018).

36. Knight H, Trewavas AJ, Knight MR. Calcium signalling in Arabidopsis thaliana responding to drought and salinity. Plant J 12, 1067–1078 (1997).

37. Liang F, Sze H. A High-Affinity Ca^2+^ Pump, ECA1, from the Endoplasmic Reticulum Is Inhibited by Cyclopiazonic Acid but Not by Thapsigargin. Plant Physiol 118, 817–825 (1998).

38. Krebs M, et al. FRET-based genetically encoded sensors allow high-resolution live cell imaging of Ca^2+^ dynamics. Plant J 69, 181–192 (2012).

39. Kalladan R, Lasky JR, Chang TZ, Sharma S, Juenger TE, Verslues PE. Natural variation identifies genes affecting drought-induced abscisic acid accumulation in Arabidopsis thaliana. Proc Natl Acad Sci U S A 114, 11536–11541 (2017).

40. Zhou Y, Wang Y, Li J, Liang J. In vivo FRET–FLIM reveals ER-specific increases in the ABA level upon environmental stresses. Plant Physiol 186, 1545–1561 (2021).

41. Wang Y, Zhou Y, Liang J. Characterization of Organellar-Specific ABA Responses during Environmental Stresses in Tobacco Cells and Arabidopsis Plants. Cells 11, (2022).

42. Wang ZP, et al. Egg cell-specific promoter-controlled CRISPR/Cas9 efficiently generates homozygous mutants for multiple target genes in Arabidopsis in a single generation. Genome Biol 16, 144 (2015).

43. Costa A, et al. Ca2+-dependent phosphoregulation of the plasma membrane Ca2+-ATPase ACA8 modulates stimulus-induced calcium signatures. J Exp Bot 68, 3215–3230 (2017).

44. Valmonte GR, Arthur K, Higgins CM, MacDiarmid RM. Calcium-Dependent Protein Kinases in Plants: Evolution, Expression and Function. Plant Cell Physiol 55, 551–569 (2014).

45. Giacometti S, Marrano CA, Bonza MC, Luoni L, Limonta M, De Michelis MI. Phosphorylation of serine residues in the N-terminus modulates the activity of ACA8, a plasma membrane Ca^2+^-ATPase of Arabidopsis thaliana. J Exp Bot 63, 1215–1224 (2012).

46. Garcia Bossi J, et al. The role of P-type IIA and P-type IIB Ca^2+^-ATPases in plant development and growth. J Exp Bot 71, 1239–1248 (2020).

47. Zhang WJ, Zhou Y, Zhang Y, Su YH, Xu T. Protein phosphorylation: A molecular switch in plant signaling. Cell Rep 42, 112729 (2023).

48. Cunningham KW, Fink GR. Calcineurin-dependent growth control in *Saccharomyces cerevisiae* mutants lacking *PMC1*, a homolog of plasma membrane Ca^2+^ ATPases. J Cell Biol 124, 351–363 (1994).

49. Bagur R, Hajnóczky G. Intracellular Ca^2+^ Sensing: Its Role in Calcium Homeostasis and Signaling. Mol Cell 66, 780–788 (2017).

50. Resentini F, Ruberti C, Grenzi M, Bonza MC, Costa A. The signatures of organellar calcium. Plant Physiol 187, 1985–2004 (2021).

51. Cheng M, et al. Lanthanum(III) triggers AtrbohD- and jasmonic acid-dependent systemic endocytosis in plants. Nat Commun 12, 4327 (2021).

52. Edel KH, Kudla J. Integration of calcium and ABA signaling. Curr Opin Plant Biol 33, 83–91 (2016).

53. Rodriguez L, et al. C2-domain abscisic acid-related proteins mediate the interaction of PYR/PYL/RCAR abscisic acid receptors with the plasma membrane and regulate abscisic acid sensitivity in Arabidopsis. Plant Cell 26, 4802–4820 (2014).

54. Diaz M, et al. Calcium-dependent oligomerization of CAR proteins at cell membrane modulates ABA signaling. Proc Natl Acad Sci USA 113, E396–405 (2016).

55. Nitsche J, et al. Structural basis for activation of plasma-membrane Ca^2+^-ATPase by calmodulin. Communi Biol 1, (2018).

56. Hwang I, Harper JF, Liang F, Sze H. Calmodulin activation of an endoplasmic reticulum-located calcium pump involves an interaction with the N-terminal autoinhibitory domain. Plant Physiol 122, 157–167 (2000).

57. Naito Y, Hino K, Bono H, Ui-Tei K. CRISPRdirect: software for designing CRISPR/Cas guide RNA with reduced off-target sites. *Bioinformatics (Oxford*, England) 31, 1120–1123 (2015).

58. Zhou Y, Wang Y, Zhang D, Liang J. Endomembrane-biased dimerization of ABCG16 and ABCG25 transporters determines their substrate selectivity in ABA-regulated plant growth and stress responses. Mol Plant 17, 478–495 (2024).

